# The loss of DNA polymerase epsilon accessory subunits POLE3-POLE4 leads to BRCA1-independent PARP inhibitor sensitivity

**DOI:** 10.1101/2023.09.21.558850

**Authors:** Hasan Mamar, Roberta Fajka-Boja, Mónika Mórocz, Eva Pinto Jurado, Siham Zentout, Alexandra Mihuț, Anna Georgina Kopasz, Mihály Mérey, Rebecca Smith, Lajos Haracska, Sébastien Huet, Gyula Timinszky

## Abstract

The clinical success of PARP1/2 inhibitors prompts the expansion of their applicability beyond homologous recombination deficiency. Here, we demonstrate that the loss of the accessory subunits of DNA polymerase epsilon, POLE3 and POLE4, sensitizes cells to PARP inhibitors. We show that the sensitivity of POLE4 knockouts is not due to a compromised response to DNA damage or homologous recombination deficiency. Instead, POLE4 deletion generates replication stress with the accumulation of single-stranded DNA gaps upon PARP inhibitor treatment. In POLE4 knockouts, replication stress leads to elevated DNA-PK signaling revealing a role of POLE4 in regulating DNA-PK activation. Moreover, POLE4 knockouts show synergistic sensitivity to the co-inhibition of ATR and PARP. Finally, POLE4 loss enhances the sensitivity of BRCA1-deficient cells to PARP inhibitors and counteracts acquired resistance consecutive to restoration of homologous recombination. Altogether, our findings establish POLE4 as a promising target to improve PARP inhibitor driven therapies and hamper acquired PARP inhibitor resistance.

## INTRODUCTION

PARP inhibitors (PARPi) emerged as a promising therapeutic approach for the treatment of cancers with mutations in the breast cancer susceptibility genes BRCA1/2 nearly two decades ago^1, 2^. BRACA1 and BRCA2 are pivotal for DNA double strand break (DSB) repair via the high-fidelity homologous recombination (HR) pathway. Mutations in the BRCA1/2 genes force the cells to rely on the error-prone non-homologous end joining (NHEJ) DSB repair, which leads to genomic instability^3^.

PARP1 is the main writer of the posttranslational modification, ADP-ribosylation in response to DNA damage^4^. PARP1 has a crucial role in the DNA damage response as it recruits rapidly to the DNA lesions modifying itself and nearby targets by adding ADP-ribose moieties on specific protein residues. The poly(ADP-ribose) (PAR) chains generated by PARP1 trigger the recruitment of chromatin remodelers and DNA repair factors involved in early steps of the DNA damage response^5, 6^.

PARPi not only inhibit ADP-ribosylation signaling but also increases PARP1 retention on sites of DNA damage causing a so-called “PARP trapping” phenomenon, which primarily underlies PARPi sensitivity^7^. These PARP1-DNA adducts are thought to be converted into DSBs during replication^8^, leading to genomic instability and increased cell death in the case of HR deficiencies observed in cells displaying a BRCAness phenotype^9^. It is this Achilles heel that is exploited in the treatment of BRCA-deficient tumors with PARPi.

Since the approval of PARPi in the clinic, extensive work has been done to expand their therapeutic spectrum beyond the BRCAness phenotype. For example, synthetic lethality with PARPi has been reported upon loss of Histone PARylation Factor 1 (HPF1)^10^, defects in the ribonucleotide excision repair pathway^11^, impairments of resolving trapped PARP1^12–15^ and loss of factors of the Fanconi anemia pathway^16^. More recently, PARPi sensitivity has been linked to the induction of single-stranded DNA (ssDNA) gaps either from unprocessed Okazaki fragments or unrestrained fork progression ultimately causing the cells to experience replication stress^17, 18^.

Unbiased knockout screens to identify genes underlying PARPi resistance suggested the loss of POLE3 and POLE4 to be synthetic lethal with PARPi^12, 19, 20^. POLE3 and POLE4 are subunits of DNA polymerase epsilon (POLε). POLε is a protein complex mainly responsible for replicating the DNA leading strand during S phase^21^. It consists of four subunits, the catalytic core composed of POLE1 along with POLE2 and the aforementioned accessory factors POLE3 and POLE4. Pol2 and Dpb2, the yeast orthologues of POLE1 and POLE2, respectively, are essential for viability but not Dpb3 (POLE4 in mammals) or Dpb4 (POLE3 in mammals)^22^. Deletion of Dpb3 and Dpb4 does not stall replication but instead, reduces the processivity of the Pol2-Dpb2 subcomplex due to unstable binding to DNA^23^. This role in stabilizing the POLε complex becomes critical upon replication stress as shown by increased sensitivity to hydroxyurea (HU) upon loss of Dpb4^24^. In addition, while Dpb3 is important for normal cell-cycle progression^25^, Dpb4 was reported to promote activation of the checkpoint kinase Mec1 (ATR in humans) upon replication stress^24^. Importantly, similar sensitivity can be observed in mice fibroblasts lacking POLE4^26^.

Both POLE3 and POLE4 have histone-fold domains and form a H2A-H2B-like heterodimer^27^ which displays H3-H4 histone chaperon activity *in vitro*^28^. More specifically, mice and yeast orthologs of POLE3 and POLE4 were shown to facilitate parental H3-H4 histone deposition on the leading strand keeping symmetrical segregation of histones between the two DNA strands^29, 30^. Consistent with their role in chromatin assembly, these accessory subunits were also shown to regulate heterochromatin silencing in budding and fission yeasts^31, 32^. Interestingly, the yeast ortholog of POLE3 (Dpb4) plays a dual role in this process depending on the complex it is part of **(Fig. 1A)**. On the one hand, as part of the POLε complex, the Dpb4-Dpb3 subcomplex ensures heterochromatin inheritance. On the other hand, within the yeast ortholog of the chromatin remodeling and chromatin-accessibility complex (CHRAC), the subcomplex Dpb4-dls1 (CHRAC15 in humans) is important for the inheritance of an expressed state^31^. As part of the CHRAC complex, Dpb4 also promotes histone removal at the vicinity of DSBs to facilitate DNA end resection^33^, while through its interaction with Dpb3 in the POLε complex, it regulates the activation of the yeast checkpoint kinase Rad53 (CHK2 in humans), which is the effector kinase of Mec1/ATR in yeast^33, 34^.

**Figure 1:**
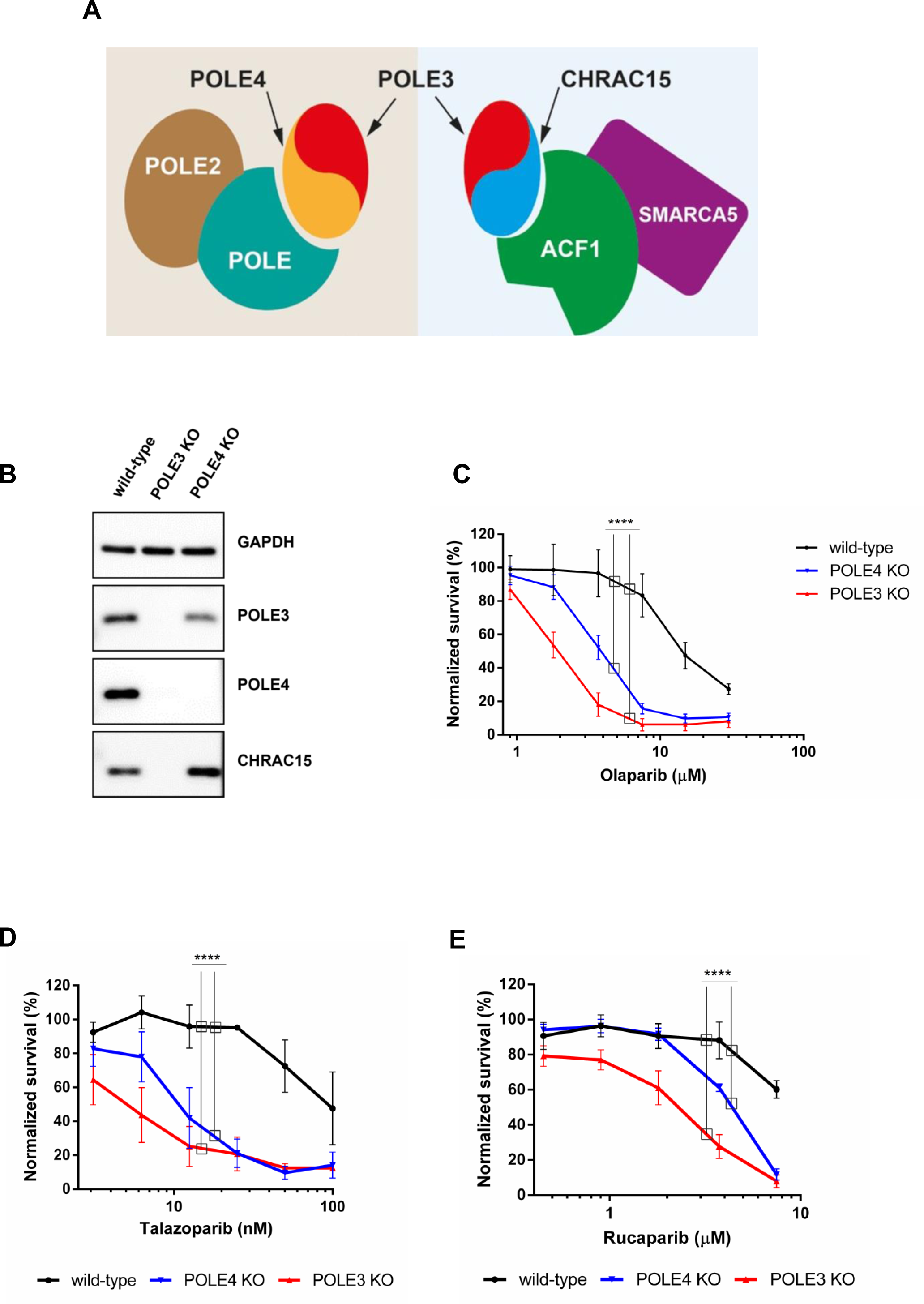
Loss of POLE3 or POLE4 induces PARPi sensitivity. (A) Schematic representation of the accessory subunits POLE3 and POLE4 within POLε and CHRAC complexes. (B) Western blot showing the levels of POLE3, POLE4 and CHRAC15 in HeLa wild-type, POLE4 KO and POLE3 KO cells. GAPDH is used as a loading control. (C-E) Cell survival assays demonstrating sensitivity of POLE3 KO and POLE4 KO to different PARPi compared to their parental HeLa wild-type. PARPi treatment was refreshed once during the 7-day long experiment. Mean ± SEM (n=3). Asterisks indicate *p*-values obtained by two-way ANOVA (**** *p*< 0.0001).

PARP activity has been implicated in most of the POLε associated functions including DNA repair, replication, and chromatin regulation. In the present work, we provide insight into the mechanisms underlying the synthetic lethality observed upon loss of POLE4 and PARP inhibition.

## RESULTS

### Loss of POLE3 or POLE4 causes PARPi sensitivity

We and others have previously identified the loss of POLE3 and POLE4 to sensitize cells to the PARP1/2 inhibitor Olaparib^12, 19, 20^. To confirm this finding, we employed CRISPR-Cas9 gene editing to generate POLE3 and POLE4 knock-outs (KO) in HeLa cells **(Fig. 1B, Supp.** Fig. 1A). As expected, all tested clones of POLE3 KO and POLE4 KO were hypersensitive to Olaparib treatment in a cell survival assay (**Fig. 1C**, **Supp.** Fig. 1B, C). Furthermore, POLE3 and POLE4 KOs were also sensitive to other PARP inhibitors, such as Talazoparib and Rucaparib, showing that the sensitivity was not limited to Olaparib **(Fig. 1D, E)**. Additionally, the PARPi sensitivity upon POLE3 and POLE4 loss was not exclusive to HeLa cells as U2OS cells knocked out for POLE3 or POLE4 showed similar sensitive phenotype to Olaparib **(Supp.** Fig. 1D, E).

The heterodimerization of POLE3 and POLE4 is essential for their stability^20^. Accordingly, the loss of either of them abolished or strongly reduced the expression of the other **(Fig. 1A, B**). POLE3 is a shared subunit between the POLε holoenzyme and the CHRAC complex^35^, where POLE3 is in a heterodimer with CHRAC15 instead of POLE4 **(Fig. 1A)**. Similar to the POLE3-POLE4 dimer, compromising the POLE3-CHRAC15 dimer by deleting POLE3 led to the loss of CHRAC15 expression **(Fig. 1B)**. Since POLE3 KO cells lack both POLE4 and CHRAC15, we decided to further characterize the consequences of PARPi treatment only in the POLE4 KO to avoid confounding phenotypes arising from the lack of both POLE3-POLE4 and POLE3-CHRAC15 heterodimers.

### PARP1 is essential for Olaparib-induced POLE4 KO sensitivity with no apparent defects in the DNA damage response

The toxicity of PARP inhibitors requires the presence of PARP1 in cells^7^, with recent reports also highlighting a requirement of PARP2^36, 37^. To investigate whether the sensitivity of POLE4 KO to PARPi was dependent on the presence of PARP1 or PARP2, we employed RNAi to deplete either or both factors. Cell survival assays demonstrated that the depletion of PARP1 alone was sufficient to rescue POLE4 KO sensitivity. Instead, depleting PARP2 neither reduced the PARPi sensitivity nor further improved survival of the POLE4 KO co-depleted for PARP1, suggesting that the Olaparib-induced sensitivity of POLE4 KO relies on the presence of PARP1, but not PARP2, consistent with PARP1 trapping on their cognate lesions **(Fig. 2A, Supp.** Fig. 2A).

**Figure 2:**
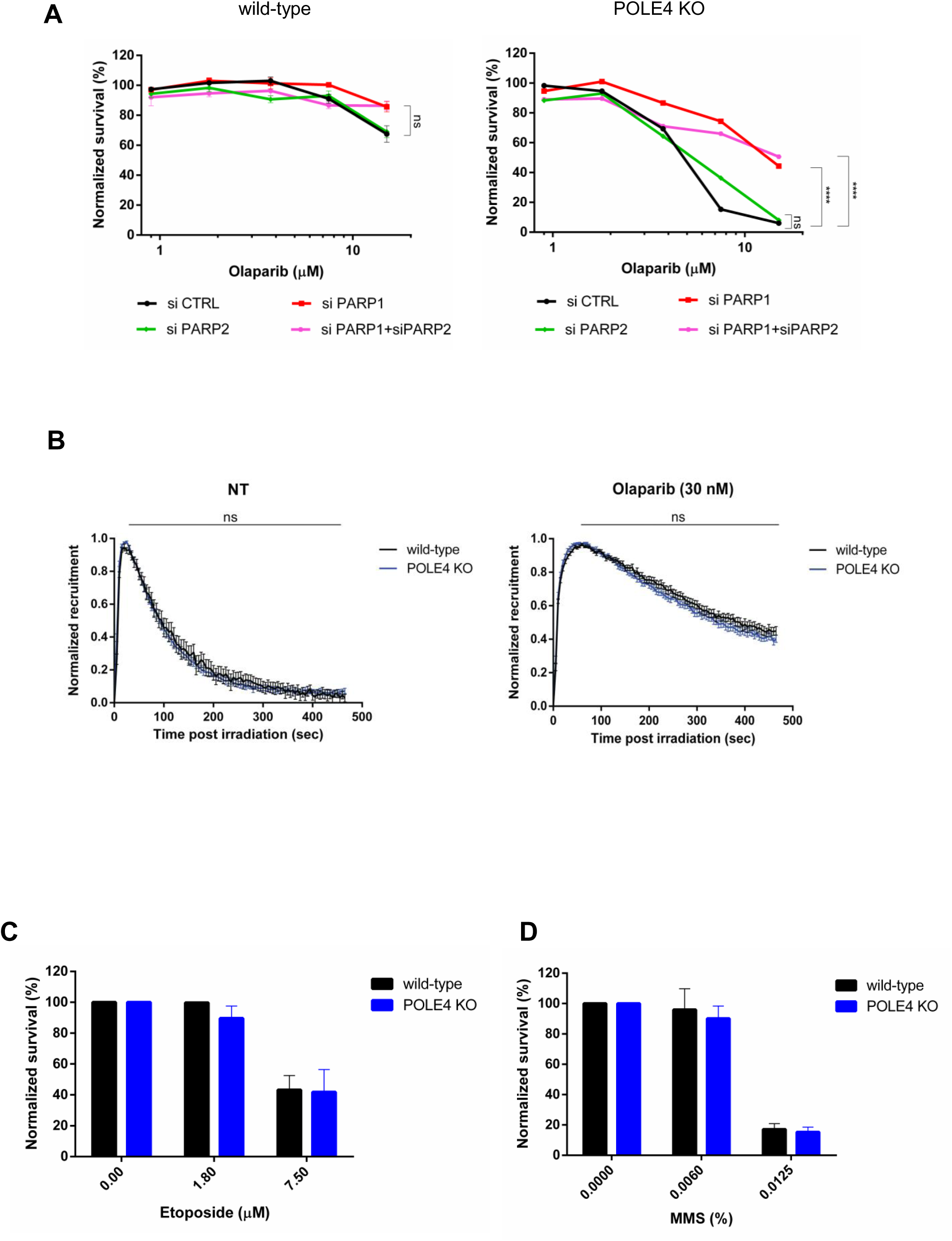
PARP1 is essential for Olaparib-induced POLE4 KO sensitivity with no apparent defects in the DNA damage response. (A) Cells survival assay showing Olaparib sensitivity of HeLa wild-type and POLE4 KO upon downregulation of PARP1, PARP2 or both of them using siRNA transfection. PARPi treatment was refreshed once during the 7-day long experiment. Mean ± SEM (n=3). Asterisks indicate *p-*values obtained by two-way ANOVA (ns. Not significant and **** *p*< 0.0001). (B) Normalized recruitment quantification of GFP-tagged PARP1 chromobody to sites of DNA damage in HeLa wild type and POLE4 KO cells in both untreated (left) or Olaparib treated (right) conditions. All data points included ± SEM. The figure is a representative experiment of three independent replicates. Measurements were analyzed using Mann-Whitney unpaired t-test. (ns. Not significant). (C, D) Cell survival of Hela wild-type and POLE4 KO cells upon treatment with etoposide (C) or MMS (D) for 1h. After the 1h treatment, cells were washed and incubated in culturing media for 7 days. Mean ± SEM (n=3).

Active ADP-ribosylation is crucial to release PARP1 from DNA. Cellular processes dampening or enhancing this signaling pathway lead to increased or reduced sensitivity to PARPi, respectively^38, 39^. Nevertheless, immunoblots showed no difference in ADP-ribose (ADPr) levels between wild-type and POLE4 KO both in the absence of genotoxic stress and after H_2_O_2_ treatment (**Supp.** Fig. 2B). These data show that loss of POLE4 is not a source of DNA lesion that leads to PARP1 activation. Moreover, it excludes POLE4 as playing a central role in the regulation of ADP-ribosylation signaling that could underlie the sensitivity of the KO cells to PARPi. Nevertheless, processes independent of ADP-ribosylation could also modulate PARP1 retention at DNA lesions^40, 41^. Thus, to more directly assess whether POLE4 regulates PARP1 mobilization from sites of DNA damage, we monitored the dynamics of endogenous PARP1 at sites of DNA damage upon laser micro-irradiation using a GFP-tagged PARP1-binding nanobody^42^. In wild-type cells, PARP1 recruited rapidly to sites of laser-induced damage before dissociating from the lesions within a time frame of few hundreds of seconds (**Fig. 2B, Supp.** Fig. 2C). As expected, this release was delayed upon Olaparib treatment **(Fig. 2B, Supp.** Fig. 2C). PARP1 kinetics at sites of laser irradiation were similar in POLE4 KO and wild-type cells, irrespective of the presence of PARPi (**Fig. 2B, Supp.** Fig. 2C) suggesting that POLE4 did not regulate PARP1 dynamics at sites of DNA damage.

The lack of apparent changes in ADPr-signaling and PARP1 dynamics at DNA lesions upon loss of POLE4 hints against a role of POLE4 in DNA repair. Consistent with this, POLE4 KO were not sensitive to genotoxic stress induced by methyl methanesulfonate (MMS) or etoposide treatments (**Fig. 2C, D**). This is in line with earlier reports that observed no sensitivity to camptothecin (CPT) or MMS in a Dpb3-deficient yeast strain^43^, or to ionizing radiation in POLE4-deficient mouse fibroblasts^26^.

Altogether these data indicate that PARPi sensitivity of POLE4 KO is not the consequence of impaired PARP1 mobilization from sites of damage or defects in the DNA damage response.

### POLE4 loss increases PARPi-induced ssDNA gaps and leads to replication stress

PARPi was reported to induce ssDNA gaps behind replication forks through the suppression of fork reversal^18^. To test whether such gaps were formed upon Olaparib treatment in cells lacking POLE4, we employed the non-denaturing BrdU immunostaining assay, where the specific antibody against BrdU is unable to bind the nucleotide analog in native conditions unless there is a single stranded DNA gap opposite it, therefore making the intensity of BrdU staining an indicator of the ssDNA gaps levels in the cell^17, 18^. With this assay, POLE4 KO cells displayed a striking increase in the intensity of BrdU staining upon Olaparib treatment compared to their wild-type counterparts, indicating a dramatic accumulation of unprocessed ssDNA gaps in these cells **(Fig. 3A, B)**.

**Figure 3:**
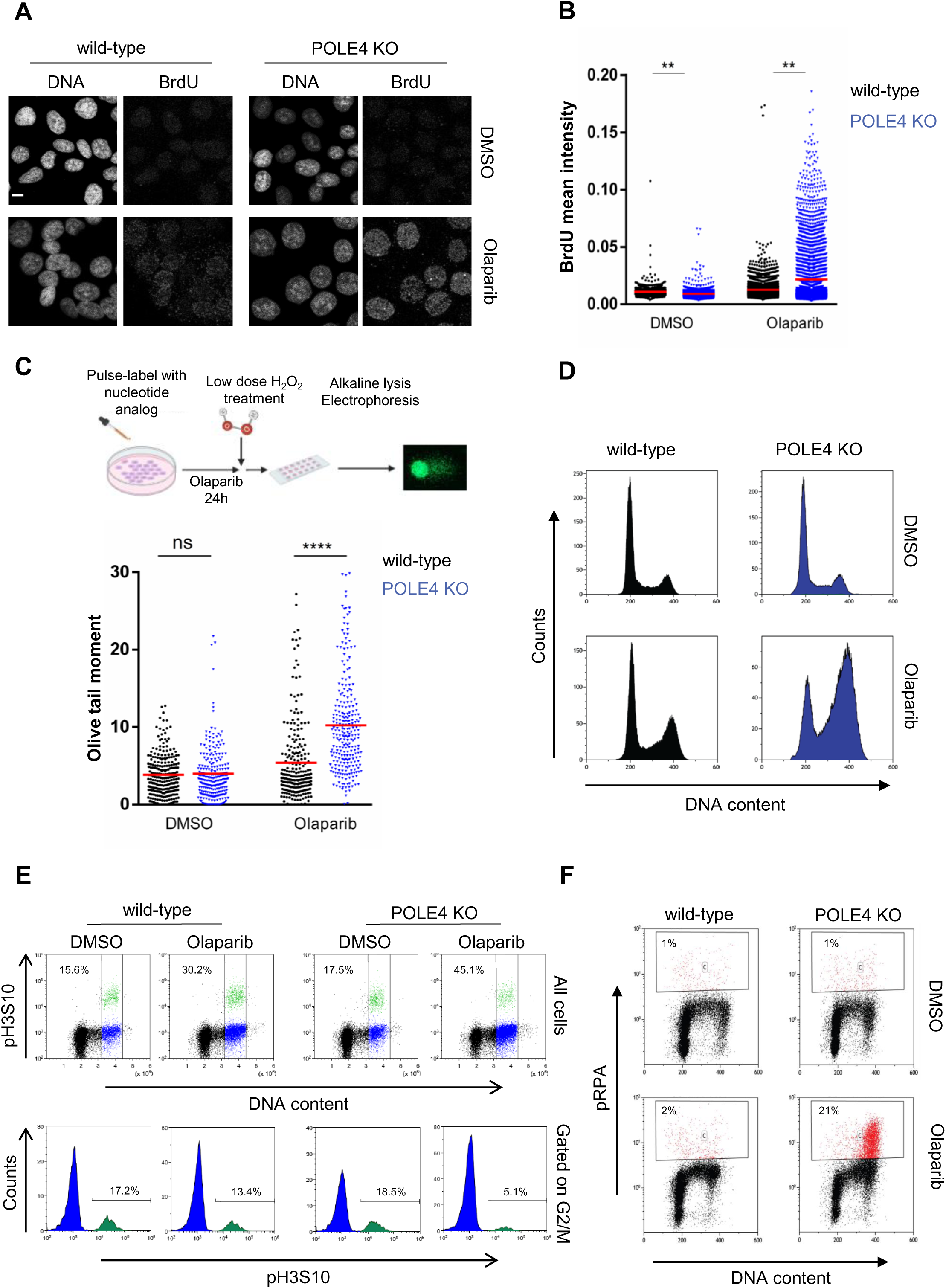
POLE4 loss increases PARPi-induced ssDNA gaps and leads to replication stress. (A, B) Immunofluorescence experiment of native BrdU staining. Cells with the indicated genotypes were incubated with BrdU (20 µM, 48h), then treated with Olaparib (10 µM, 24h) or the control vehicle DMSO. (A) Representative images are shown. Scale bar, 10 μm. (B) Mean BrdU intensity of all scored cells was blotted. The graph represents one experiment out of three independent repetitions. Asterisks indicate *p*-values obtained by one-way ANOVA (** *p*< 0.01). (C) (Top) Schematic of BrdU-Comet experiment. (Bottom) quantification of olive tail moment of HeLa wild-type and POLE4 KO cells treated with Olaparib (20 µM, 24h) or DMSO. The figure is a representative of three independent experiments. Asterisks indicate *p*-values obtained by one-way ANOVA (ns. Not significant, **** *p*< 0.0001). (D) Representative FACS experiment showing cell-cycle profile of cells with the indicated genotypes with or without Olaparib treatment (5 μM, 24 h). (E) (Top) Representative FACS experiment for distinguishing the mitotic cells by positive staining of pH3S10 (green) from G2 phase cells (blue) in Hela wild-type or POLE4 KO, treated or not with Olaparib (5 μM, 24 h). Percentages of cells in G2/M relative to all scored cells are shown. (Bottom) Histograms showing the cells gated on G2/M from top panel. Numbers represent the percentage of mitotic cells within the G2/M population. (F) Flow cytometry of Hela wild-type or POLE4 KO cells after culturing for 24h with Olaparib (5 μM) or DMSO. The cells were fixed and stained with anti-pRPA and propidium-iodide (DNA content). Percentages of pRPA positive cells (red) are shown. The figure is a representative of three independent experiments.

To further confirm this observation, we performed a variation of a comet assay, as so called BrdU comet assay, where pulse labeling the cells with a nucleotide analog was used to highlight newly synthesized DNA, rendering the assay S-phase specific^44, 45^. The assay was further modified by application of a short, low dose of H_2_O_2_ treatment immediately prior to embedding the cells in agarose, to convert single stranded gaps into double stranded breaks, the resulting fragmented DNA fraction thus migrating into the comet tail^45^. Consistent with the non-denaturing BrdU staining results **(Fig. 3A, B)**, we observed a significant increase in the Olive tail moment of POLE4 KO cells treated with Olaparib, underscoring the higher levels of ssDNA gaps in these cells due to PARPi treatment **(Fig. 3C, Supp.** Fig. 3A).

Sensitivity to PARPi has been linked to defects in Okazaki fragment processing during DNA replication^17^, and ADPr signal was reported to correlate with the amount of unligated Okazaki fragments^46^. Importantly, detection of significant ADPr signal was only possible upon inhibition of the poly(ADP-ribose) glycohydrolase (PARG)^46^. To test whether POLE4 had a role in Okazaki fragment processing, we loaded POLE4 KO cells with the amine-reactive carboxyfluorescein diacetate, succinimidyl ester (CFSE) cell tracker dye and mixed it with its wild-type unlabeled cells to ensure each cell line undergoes the exact same conditions when performing the experiment and imaging. Following the mixing of cells, we assessed PAR levels in replicating cells treated with PARGi. The lack of difference in ADPr levels between POLE4 KO and wild-type cells suggested similar amounts of Okazaki fragments in both cell lines (**Supp.** Fig. 3B, C). Co-inhibition of PARG and Fen1, an enzyme responsible for processing Okazaki fragments^46^, increased further the ADPr signal compared to PARG inhibition alone but again to comparable levels in both wild-type and POLE4 KO (**Supp.** Fig. 3B, C), indicating that POLE4 loss neither alters the processing of Okazaki fragments nor leads to increased S-phase specific PARP activity.

Accumulation of unprocessed ssDNA gaps induces replication stress and alters cell cycle progression^47^. Notably, PARPi treatment was associated with the accumulation of cells in late S and G2/M phase, which was much more prominent in POLE4 KO cells **(Fig. 3D)**. Yet, there were no major differences in cell cycle progression between wild-type and POLE4 KO cells in untreated conditions **(Fig. 3D)** supporting the notion that it is only upon PARPi treatment that the loss of POLE4 impacts replication. To refine whether Olaparib leads to G2 or mitotic arrest, we quantified a mitotic marker, histone H3 phosphorylated on Ser10 (pH3S10), together with propidium iodide by flow cytometry **(Fig 3E)**. Despite the large accumulation of cells with DNA content characteristic to G2/M detected in Olaparib-treated samples **(Fig. 3E upper row)**, the ratio of mitotic cells within this population decreased, especially in POLE4 KO **(Fig. 3E lower row)**. This suggests that POLE4 KO cells are prevented from entering mitosis upon PARP inhibition, presumably due to the accumulation of ssDNA gaps. By examining the cell cycle dependent phosphorylation on residue T21 of the ssDNA binder replication protein A subunit (pRPA) by flow cytometry, we did not observe an enhanced signal in wild-type cells either treated or not with Olaparib **(Fig. 3F)**. In sharp contrast, POLE4 KO cells showed strong pRPA positivity in G2/M phase of the cell cycle upon Olaparib treatment suggesting an induction of replication stress **(Fig. 3F)**. Consistent with a role of POLE4 in preventing ssDNA gaps in the presence of replicative stress, POLE4 KO were hypersensitive to replication stress induced by other agents as well, such as HU or Ataxia-telangiectasia mutated and RAD3-related (ATR) inhibitor (ATRi) treatment in cell survival assays **(Supp.** Fig. 3D, E), in line with previous reports^20, 48, 49^.

Taken together, these results demonstrate that the loss of POLE4 does not increase ssDNA gaps or unprocessed Okazaki fragments when ADP-ribosylation signaling is active, rather, POLE4 is essential to suppress PARPi-induced ssDNA gaps and subsequent replication stress induction.

### PARPi-induced replication stress in POLE4 KO cells is controlled by PIKKs

During DNA replication, the Phosphatidylinositol 3-kinase-related kinase (PIKK) family member ATR orchestrates origin firing, protects replication forks and regulates cell cycle progression^50^. ATRi can induce ssDNA accumulation, which can lead to phosphorylation and activation of other PIKKs, such as DNA-dependent protein kinase (DNA-PK) and Ataxia-telangiectasia mutated (ATM)^50^. As POLE4 KO cells were sensitive to replication stress **(Supp.** Fig. 3D, E), we sought to further examine replication stress signaling in these cells. While DNA-PK phosphorylation was not enhanced in wild-type cells upon Olaparib and/or ATRi treatment, it was markedly elevated in POLE4 KO after 24 hours of ATRi treatment alone or combined with Olaparib. **(Fig. 4A)**. Phosphorylated ATM (pATM) – an indicator of ATM activation – was modestly elevated upon incubation of the wild-type cells either with Olaparib or with ATRi alone, while strongly enhanced in the presence of their combination. This was further increased in POLE4 KO cells both in the case of single Olaparib or ATRi treatment and when combined, indicating stronger activation of ATM when POLE4 is missing (**Fig. 4A).** These results suggest an interplay of PIKKs in POLE4 KO cells in response to replication stress.

**Figure 4:**
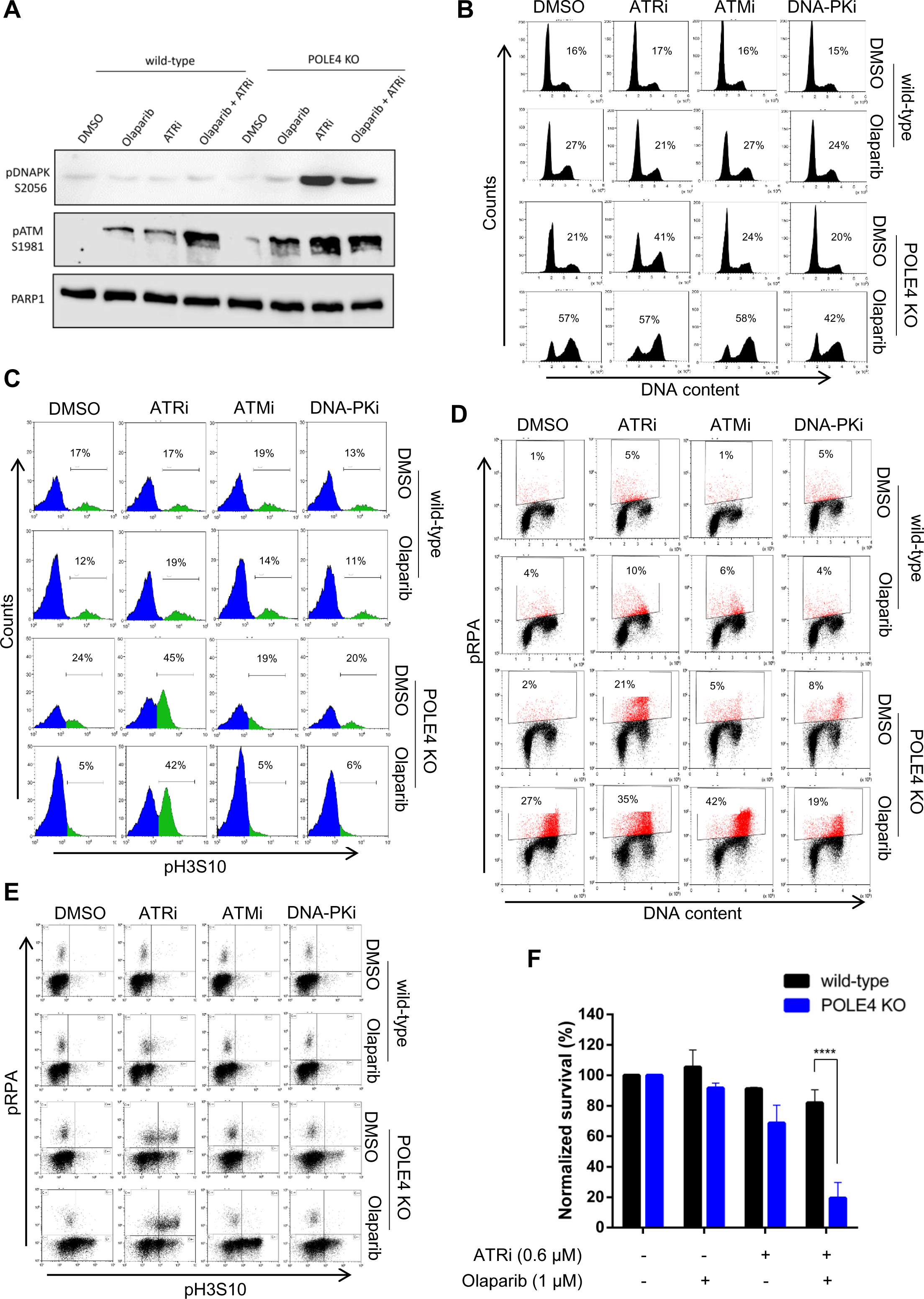
PARPi-induced replication stress in POLE4 KO cells is controlled by PIKKs. (A) Western blot of HeLa wild-type and POLE4 KO cells with the indicated antibodies upon treatment with Olaparib (5 µM, 24h), ATRi (5 µM, 24h) or both of them. PARP1 is used as a loading control. (B) Representative FACS experiment showing cell-cycle profile of HeLa wild-type and POLE4 KO cells after 24h of Olaparib (5 μM) and/or ATRi (5 μM), ATMi (5 μM) DNA-PKi (5 μM) treatment. DMSO was used as a solvent control. Numbers represent the percentages of G2/M population relative to all cells. (C) Flow cytometry of Hela wild-type or POLE4 KO cells treated for 24h with Olaparib (5 μM) and/or ATRi (5 μM), ATMi (5 μM), DNA-PKi (5 μM) and then stained with anti-pH3S10 and propidium-iodide. Histograms show the cells gated on G2/M according to DNA content, and the ratio of mitotic cells was determined by pH3S10 positivity (green). Percentages of pH3S10 positive cells relative to the population of G2/M are shown. DMSO was used as a solvent control. (D) Flow cytometry of Hela wild-type or POLE4 KO cells after 24h of Olaparib (5 μM) and/or ATRi (5 μM), ATMi (5 μM), DNA-PKi (5 μM) treatment. DMSO was used as a solvent control. The cells were fixed and stained with anti-pRPA and propidium-iodide (DNA content). Percentages of pRPA positive cells (red) are shown. The figure is a representative of three independent experiments. (E) Flow cytometry of Hela wild-type or POLE4 KO cells after 24h of Olaparib (5 μM) and/or ATRi (5 μM), ATMi (5 μM), DNA-PKi (5 μM) treatment. DMSO was used as a solvent control. The cells were fixed and stained with anti-pH3S10 and anti-pRPA. (F) Cell survival assay of HeLa wild-type and POLE4 KO cells. The columns represent normalized survival of the cells upon the indicated treatments. The treatment was refreshed once during the 7-day experiment. Mean ± SEM (n=3). Asterisks indicate *p-*values obtained by two-way ANOVA (**** *p*< 0.0001).

Therefore, we assessed if the cell cycle profile of wild-type and POLE4 KO cells was modified by PIKK inhibitors combined with Olaparib treatment. The cell cycle of wild-type cells was not affected by ATRi, ATMi or DNA-PKi alone, but there was a G2/M accumulation in Olaparib treated samples with or without PIKK inhibitors **(Fig. 4B)**. In line with the ATRi sensitivity of POLE4 KO, ATRi treatment alone, but not other PIKK inhibitors, caused marked accumulation of G2/M cells in POLE4 KO **(Fig. 4B).** ATR and ATM inhibition had little effect on the Olaparib-induced cell cycle arrest, in contrast to DNA-PKi, which slightly reduced it **(Fig. 4B)**. When distinguishing G2 and M phase cells with the mitotic marker pH3S10, ATR inhibition promoted the transition of POLE4 KO cells into mitosis, even if PARP1 was inhibited, which otherwise blocked them in G2 phase. In contrast, the proportion of mitotic cells did not change in the presence of other PIKK inhibitors (**Fig. 4C).** These data establish ATR as the major checkpoint kinase responsible for cell cycle arrest at the G2/M transition in POLE4 KO.

Next, we aimed to address which PIKK was responsible for the pRPA signal upon Olaparib treatment. Although Olaparib treatment caused cell cycle arrest both in wild-type and POLE4 KO cells, we detected marked pPRA signal only in cells lacking POLE4 **(Fig. 3F and Fig. 4D).** Treatment with PIKK inhibitors did not significantly alter the percentage of pRPA positive wild-type cells upon Olaparib treatment. In contrast, ATRi alone induced strong RPA phosphorylation in POLE4 KO cells (**Fig. 4D**). Compared to this, the pRPA signal with ATMi or DNA-PKi treatment alone was only marginally increased in POLE4 KO. On the other hand, PIKK inhibitors modified the stress response of POLE4 KO to Olaparib: ATRi and ATMi enhanced the percentage of pRPA positive cells, while DNA-PKi decreased it (**Fig. 4D**). Notably, DNA-PK inhibition also alleviated the ATRi-induced cell cycle arrest and RPA phosphorylation in POLE4 KO **(Supp.** Fig. 4A, B). These results point towards a role of DNA-PK in promoting RPA phosphorylation and subsequent stress signaling. This is in line with the western blotting data, where ATRi induced phosphorylation of DNA-PK in POLE4 KO cells (**Fig. 4A**), ultimately leading to RPA phosphorylation.

Interestingly, the double staining of pH3S10 and pRPA revealed that Olaparib-induced pRPA positive POLE4 KO cells are prevented from entering mitosis, unless they were released from the control of ATR **(Fig. 4E)**. ATR inhibition therefore forces the cells into premature mitosis, which could finally lead to replication catastrophe and reduced cell survival^48, 51^. To address this possibility, we performed cell survival assays using a low concentration of ATRi alone or combined with Olaparib. While single treatment with low dose of either Olaparib or ATRi had minor effect on the survival of POLE4 KO cells, combining both synergistically killed POLE4 KO cells **(Fig. 4F)**.

These results emphasize the importance of ATR signaling in restraining POLE4 KO from entering mitosis upon PARPi-induced replication stress. They also establish the loss of POLE4 as a major sensitizing event to co-treatment with ATRi and PARPi, a combination that is being tested in clinical trials^52^. Furthermore, these data suggest that POLE4 plays a role in suppressing DNA-PK signaling in response to replication stress, which is most obvious in response to ATRi.

### POLE4 acts parallel to BRCA1 in inducing PARP inhibitor sensitivity

Since PARPi sensitivity was first described in BRCA-deficient cells displaying impaired HR^1, 2^, we aimed to check whether PARPi-induced replication stress response could be detected when BRCA1 was missing. Similar to POLE4 KO, downregulating BRCA1 resulted in increased levels of pRPA upon Olaparib treatment (**Fig. 5A**). Strikingly, co-depletion of POLE4 and BRCA1 had a synthetic impact on pRPA levels compared to single depletion **(Fig. 5A)**. This suggests that POLE4 might function parallel to BRCA1, and that it is not part of the canonical HR pathway.

**Figure 5:**
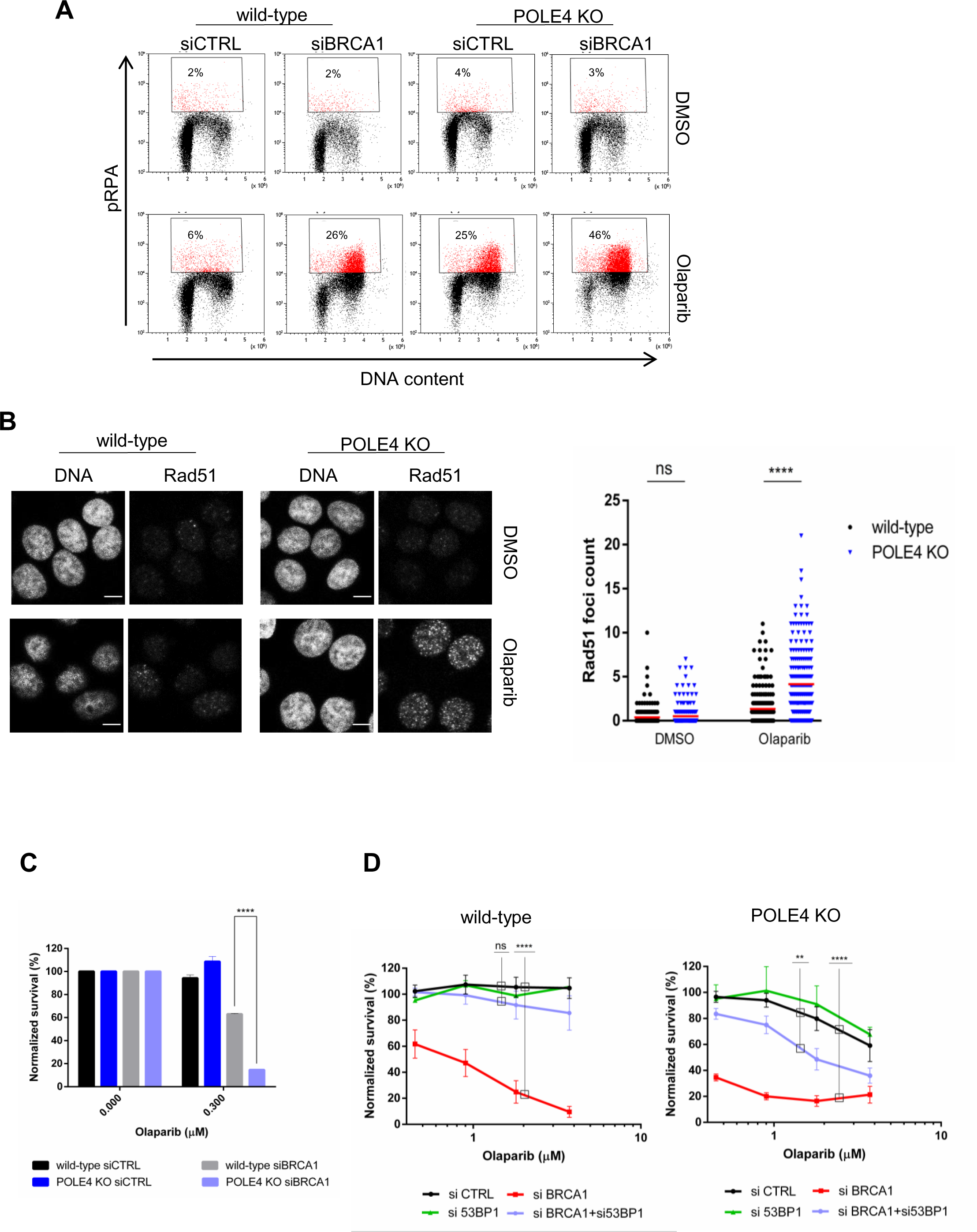
POLE4 acts parallel to BRCA1 in inducing PARP inhibitor sensitivity. (A) Flow cytometry of Hela wild-type or POLE4 KO cells with downregulated BRCA1, treated with Olaparib (5 μM, 24 h) or DMSO. The cells were fixed and stained with anti-pRPA and propidium-iodide (DNA content). Percentages of pRPA positive cells (red) are shown. The figure is a representative of three independent experiments. (B) (Left) Representative images of immunofluorescence experiment of Rad51 foci formation in HeLa wild-type and POLE4 KO cells upon treatment of Olaparib (10 µM, 48h), Scale bar, 10 μm. (Right) Quantification of Rad51 foci count in the indicated cell lines upon the indicated treatment. The experiment is representative of three independent repetitions. Asterisks indicate *p*-values obtained by one-way ANOVA (ns. Not significant, **** *p*< 0.0001). (C) Cell survival assay of HeLa wild-type and POLE4 KO cells transfected with the indicated siRNA and treated with the indicated concentration of Olaparib for 7 days, with the treatment being changed once, before calculating the relative survival normalized to the untreated samples of each genotype. Mean ± SEM (n=3). Asterisks indicate *p*-values obtained by two-way ANOVA (**** *p*< 0.0001). (D) Cell survival assay of HeLa wild-type and POLE4 KO cells downregulated of either BRCA1, 53BP1 or both of them using siRNA transfection and treated with the indicated concentrations of Olaparib for 7 days, with the treatment being changed once. Mean ± SEM (n=3). Asterisks indicate *p*-values obtained by two-way ANOVA (ns. Not significant, ** *p*< 0.01, **** *p*< 0.0001).

To confirm this hypothesis, we examined PARPi-induced Rad51 foci formation by confocal microscopy. The recombinase Rad51 is a crucial protein in the process of HR: following DNA end-resection, Rad51 binds ssDNA overhangs and leads the homology search and strand invasion to facilitate homology-directed repair^1, 2^. Consistent with previous reports describing impairment in HR, BRCA1-deficient cells displayed reduced Rad51 foci formation compared to the BRCA1 proficient controls (**Supp.** Fig. 5A, B). Conversely, POLE4 KO cells were able to efficiently form Rad51 foci upon Olaparib treatment, even to a higher extent than their wild-type counterpart (**Fig. 5B)**. This observation can be attributed to the elevation of ssDNA gaps we described previously in POLE4 KO following PARPi (**Fig. 3A, B**).

Since POLE4 is not redundant in function with BRCA1, we reasoned that PARPi sensitivity could be potentiated if both proteins were missing. To that end, we downregulated BRCA1 in wild-type and POLE4 KO and challenged the cells with a low dose of Olaparib. BRCA1 depletion in POLE4 KO cells resulted in massive killing of these cells in comparison to the loss of either POLE4 or BRCA1 alone (**Fig. 5C**), indicating that POLE4 might serve as a potential target for enhancing sensitivity of BRCA1-deficient tumors to PARPi.

A common mechanism for BRCA1-deficient tumors acquiring resistance to PARPi is the rewiring of HR through loss of the NHEJ factor 53BP1^53–55^. Given that sensitivity of POLE4 KO to PARPi is not going through defective HR, we sought to investigate whether targeting POLE4 could overcome PARPi resistance observed upon loss of 53BP1 in BRCA1-deficient cells^53–55^. To that end, we utilized RNAi-mediated downregulation of either BRCA1, 53BP1 or their combination in wild-type and POLE4 KO cells. Consistent with previous reports, downregulating BRCA1 in wild-type cells sensitized them to Olaparib, which was rescued with combined depletion of BRCA1/53BP1 (**Fig. 5D, Supp.** Fig. 5C). As mentioned earlier, BRCA1 depletion in POLE4 KO cells resulted in severe sensitization to Olaparib in comparison to missing either POLE4 or BRCA1 alone **(Fig. 5D).** Significantly, the co-depletion of BRCA1/53BP1 did not rescue PARPi sensitivity of POLE4 KO as in the case of wild-type cells (**Fig. 5D, Supp.** Fig. 5C), indicating that targeting POLE4 not only enhanced PARPi synthetic lethality in BRCA1-depleted cells but also bypassed the synthetic viability induced by reactivation of HR upon 53BP1 loss in BRCA1-compromised cells^54^.

## DISCUSSION

Our results validate that POLE4 deficiency causes PARPi sensitivity. While Olaparib inhibits both PARP1 and PARP2^7^, our results provide support that the toxicity of PARP inhibitors in POLE4 KO is dependent on PARP1 rather than of PARP2, justifying the efforts of developing PARP1 specific inhibitors^56, 57^.

PARPi cytotoxicity has been linked to the accumulation of ssDNA gaps^17, 18^. Such ssDNA gaps can either be (1) the consequence of defective Okazaki fragment processing^17, 46^ or (2) due to PARPi treatment generating ssDNA gaps behind replication fork that persist due to trapped PARP1^18^. The loss of POLE4 caused severe replication stress and cell cycle arrest upon PARP inhibition. One can envision two scenarios: POLE4 could impair replication causing replication stress, which may lead to an increased requirement for PARP activity, alternatively, POLE4 could play a role in avoiding replication stress, through the regulation of intra-S-phase signaling or facilitating replication in challenging chromatin environments.

Unligated Okazaki fragments have been proposed to be a major source of PARP activity in S-phase. If POLE4 functions to ensure timely processing of Okazaki fragment, then its loss is expected to cause accumulation of ssDNA fragments even without PARPi treatment, such accumulation will be translated into increased PAR levels in S-phase cells just as in BRCA-deficient cells^17^. However, POLE4 KO cells do not show increased S-phase PAR signal compared to their wild-type counterparts upon treatment with either PARGi or the combination of PARGi and Fen1i, the latter interfering with Okazaki fragment processing underscoring that POLE4 does not participate in ligating Okazaki fragments. Furthermore, POLE4 KO have normal cell cycle and no detectable increase in replication stress signaling – unless challenged with inducers of replication stress. Therefore, a more likely scenario than POLE4 impairing replication upon its loss, is that the accelerated fork speed upon PARP inhibition^58^ ultimately leads to replication stress and the accumulation of ssDNA gaps when POLE4 is missing. This, however, raises the hypothesis that toxic PARP1 trapping is the consequence of replication stress induced ssDNA gap formation rather than trapped PARP1 being the primary source of replication stress.

Our results are more consistent with a role of POLE3-POLE4 in the replication stress response. Yeast studies identify a role of the POLε complex in the activation of S-phase checkpoint either through the C-terminal of the catalytic subunit^59^ or the accessory subunit Dpb4^24^. In line with our data, this role is observed only in response to replication stress^24, 59, 60^. Several reports have shown that the loss of POLE4 causes reduced replication origin activation in mice and worms^26, 61, 62^. These replication defects remain, however, relatively mild unless these cells are subjected to replication stress inducers^26^. These findings, together with our data that POLE4 KO cells are hypersensitive to replication stress, as well as other previous reports^20, 48, 49^, all point towards a key role of POLE4 in replication stress tolerance. Given that PARPi leads to accelerated replication fork speed^58^, it is tempting to speculate that such accelerated forks could be sufficient to trigger the accumulation of ssDNA gaps and the subsequent replication stress phenotype in cells deficient of POLE4.

DNA-PK signaling is overactivated in response to ATRi in POLE4 KO cells. The overactivation of DNA-PK signaling has been observed in HR-deficient cells upon PARPi treatment^63^, as well as in the absence of the histone chaperone ASF1 or the chromatin assembly factor CAF1 in response to DSB^64^ hinting to a possible link between chromatin structure and the regulation of DNA-PK activity. ATM activity is also elevated in POLE4 KOs as compared to wild-type cells in response to both PARPi and ATRi, and furthermore, it appears to be part of the canonical response as the inhibition of ATM in addition to PARPi further increases replication stress induced RPA phosphorylation.

The ATR kinase is critical to protect against replication stress^65^. When ssDNA accumulates, RPA protein complex binds these structures with high affinity promoting the recruitment of ATR^66^. Upon recruitment, ATR becomes active and starts a cascade of signaling events aiming to regulate several processes including DNA replication and cell-cycle progression^67, 68^. PARPi induced ssDNA gaps require an intact ATR pathway to engage as a salvage pathway to protect cells from replication stress^18^. Upon responding to these lesions, ATR will block cells from entering mitosis with unrepaired damage and reduce the replication rate to prevent potentiating replication stress^69^. Along with this, several studies have shown that combining PARPi and ATRi synergistically kills BRCA 1/2 deficient cells by causing premature mitotic release^51, 70^. One obstacle for combination therapy is the enhanced side effects caused by combining two or more drugs. This can be avoided by identifying genetic alterations that enhance susceptibility towards these drugs^48^. Here we report that POLE4 can serve as a target to enhance the sensitivity of cancer cells to the combination of PARPi and ATRi. We propose that the synergistic killing of POLE4 KO cells with this drug combination could be attributed to two main scenarios. First, PARPi-induced replication stress in POLE4 KO cells could be further enhanced when combined with ATRi, similar to what was observed upon loss of RNase H2^48, 71^. Second, ATR inhibition could overcome the G2/M block following PARPi-induced replication stress, leading to premature entry of POLE4 KO cells into mitosis **(Fig.6A).** This is similar to what has been reported with deficiency of BRCA1/2^51, 70^ or ATM^72, 73^.

**Figure 6:**
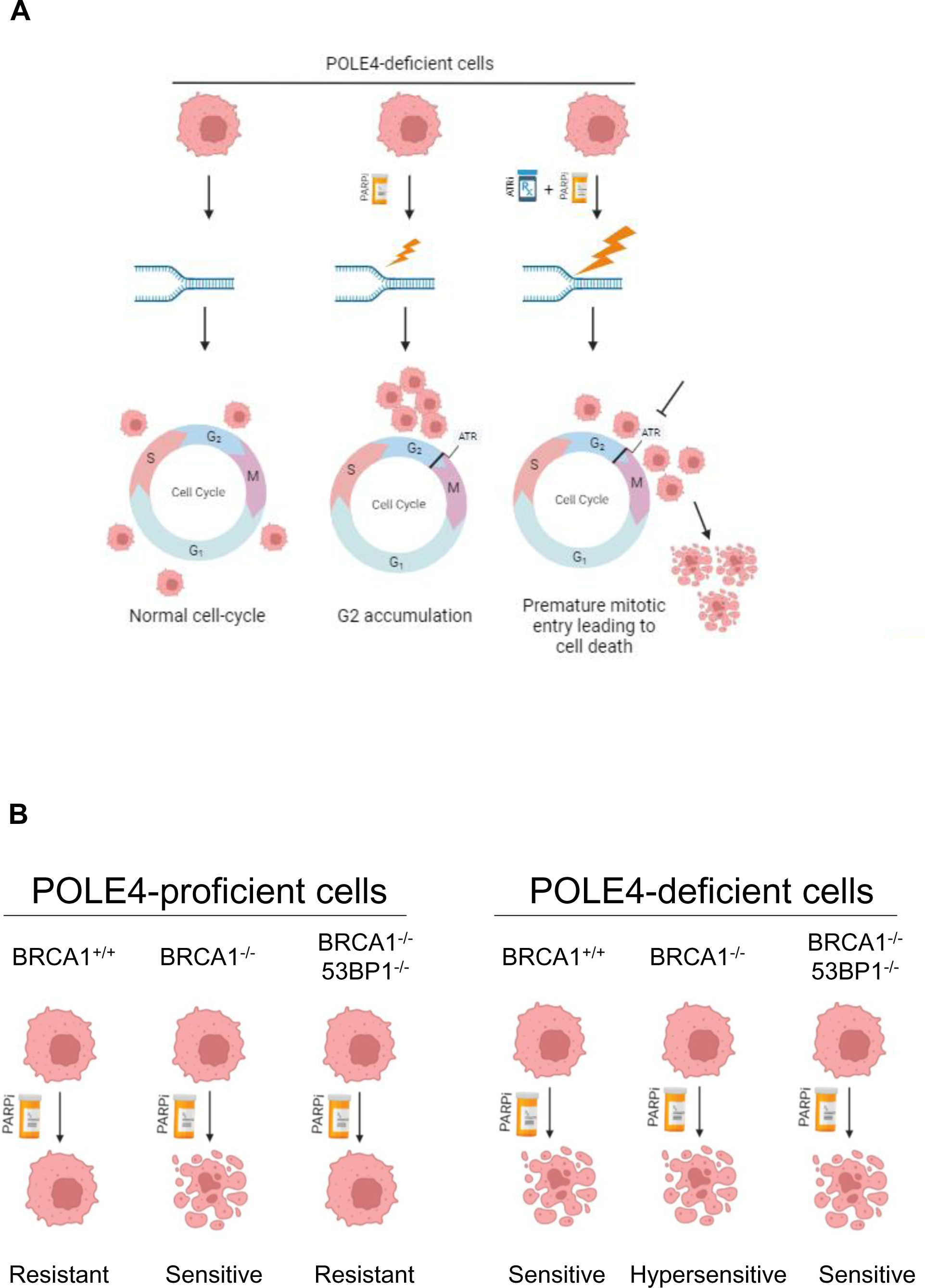
Suggested model. (A) POLE4-deficient cells have normal cell cycle progression without exogenous stress. Upon PARPi treatment, POLE4 KO show signs of replication stress that lead to G2 accumulation due to ATR checkpoint activity. Co-inhibition of PARP and ATR potentiates the replication stress in POLE4 KO and causes premature mitotic entry ultimately resulting in synergistic killing of these cells. (B) In POLE4-proficient cells, PARPi is synthetic lethal with BRCA1 deficiency, which is reversed upon loss of both BRCA1 and 53BP1 leading to PARPi acquired resistance. In POLE4-deficient background, the cells become sensitive to PARPi and this sensitivity is further enhanced upon loss of BRCA1. Importantly, the acquired resistance to PARPi due to co-depletion of BRCA1 and 53BP1 can be bypassed in POLE4-deficeint cells, highlighting a potential therapeutic exploitation in the clinic.

Moreover, in contrast to BRCA1-deficient cells, sensitivity of POLE4 KO cells to PARPi is not rescued by the restoration of HR upon 53BP1 depletion (**Fig. 6B**). Sensitivity of BRCA1-deficient tumors to PARPi can be attributed to three main mechanisms: (1) HR deficiency, (2) loss of replication fork protection, (3) defects in Okazaki fragments processing. POLE4 KO cells differ from BRCA1-deficiency in all these mechanisms, placing POLE4 in a BRCA1-independent pathway underlying PARPi resistance. Genetic deletions of POLE4 were in fact reported in cases of malignant mesothelioma^74^ and non–small cell lung cancer^75^. Therefore, our data suggest that POLE4 might serve as a biomarker for identifying tumors that can respond to PARPi treatment regardless of their HR status.

## METHODS

### Cell lines and cell culture

Cell lines used in this study were cultured in DMEM (Biosera) supplemented with 10% FBS, 100 µg/ml penicillin, 100 U/mL streptomycin and 1% NEAA and maintained at 37°C in a 5% CO2 incubator unless otherwise stated. RPE-1 p53 KO and RPE-1 p53/BRCA1 double KO were kindly gifted from Alan D. D’Andrea lab^76^ and were grown using DMEM-F12 (Biosera) supplemented with 10% FBS, 100 µg/ml penicillin, 100 U/mL streptomycin.

POLE3 KO and POLE4 KO were generated in this study from either wild-type HeLa cells or wild-type U2OS-FlpIn cells kindly provided by Ivan Ahel’s lab using CRISPR/Cas9 technology. The sgRNA sequences targeting either POLE3 or POLE4 are:

sgPOLE3: 5’-GTACAGCACGAAGACGCTGG-3’

sgPOLE4: 5’-GTCGGGATCTGCCTTCACCA-3’

### RNA Interference and Plasmid Transfection

pSpCas9(BB)-2A-Puro (PX459) V2.0 used to generate the knockouts of this study was a gift from Feng Zhang lab (Addgene, plasmid #62988)^77^. Plasmid transfections were performed using **Xfect** (Takara) according to the manufacturer’s protocol.

RNA interference experiments with siRNA (sequences in Table 1) were conducted using Dharmafect (Dharmacon) or RNAiMAX (Lipofectamine) transfection reagents according to the manufacturers’ instructions. Downregulation was verified by western blotting using specific antibodies (detailed in Table 2).

**Table1.**
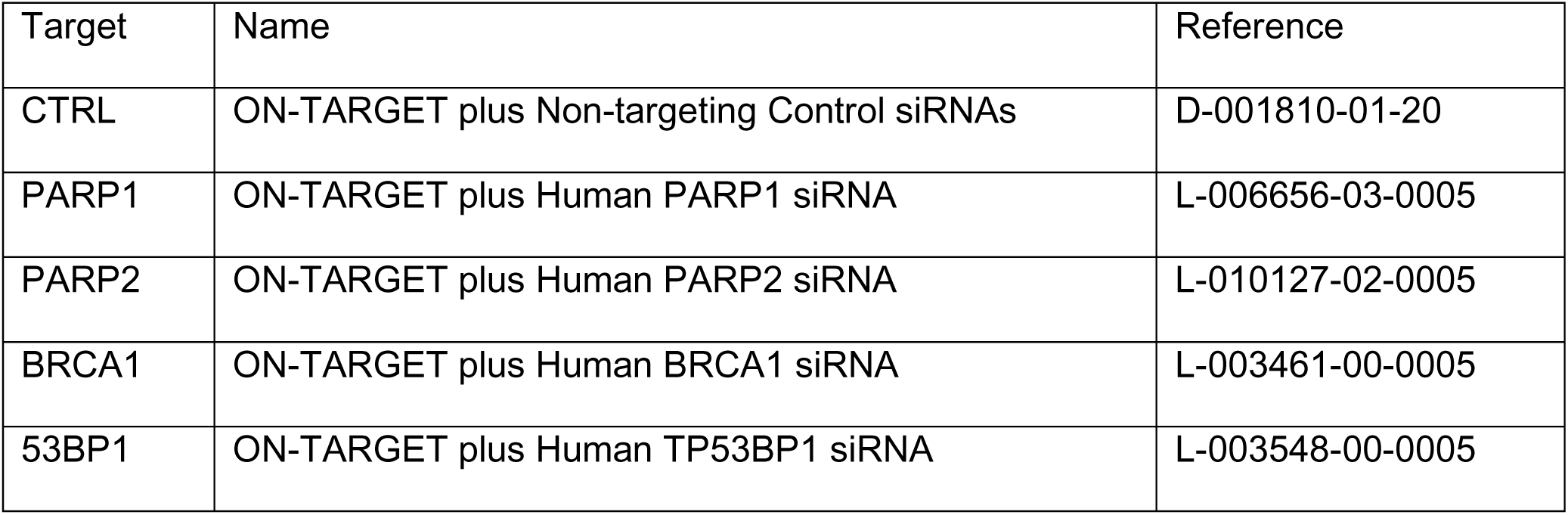
Dharmacon smart pool siRNA.

**Table 2.**
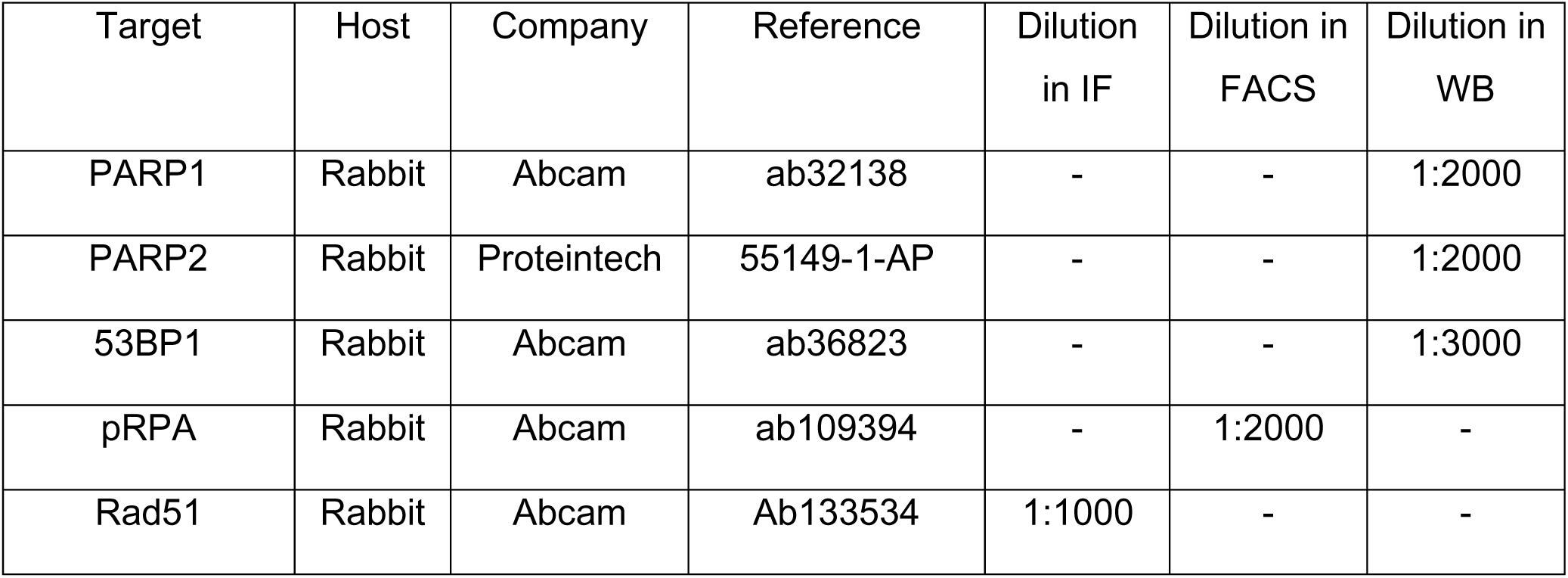
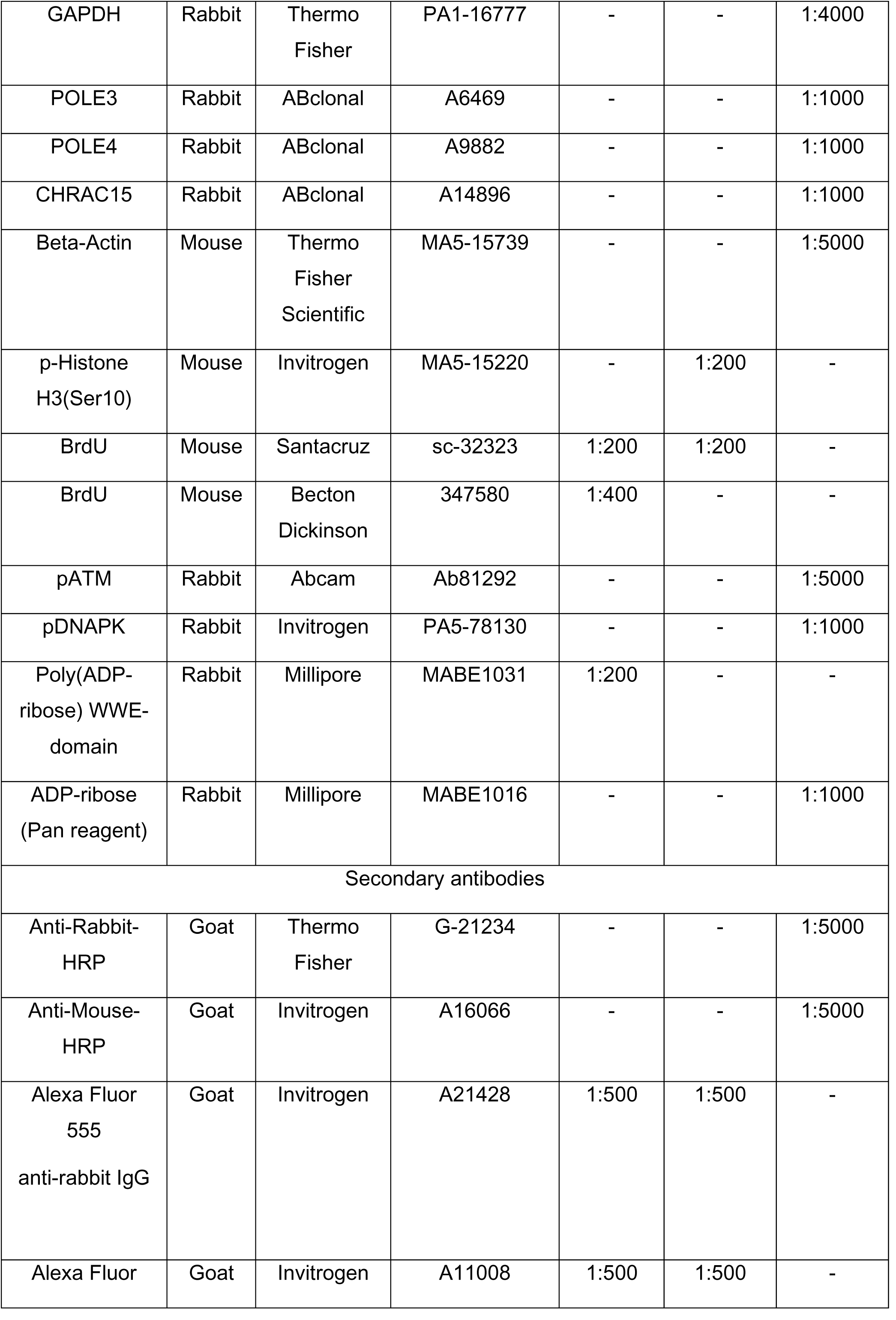
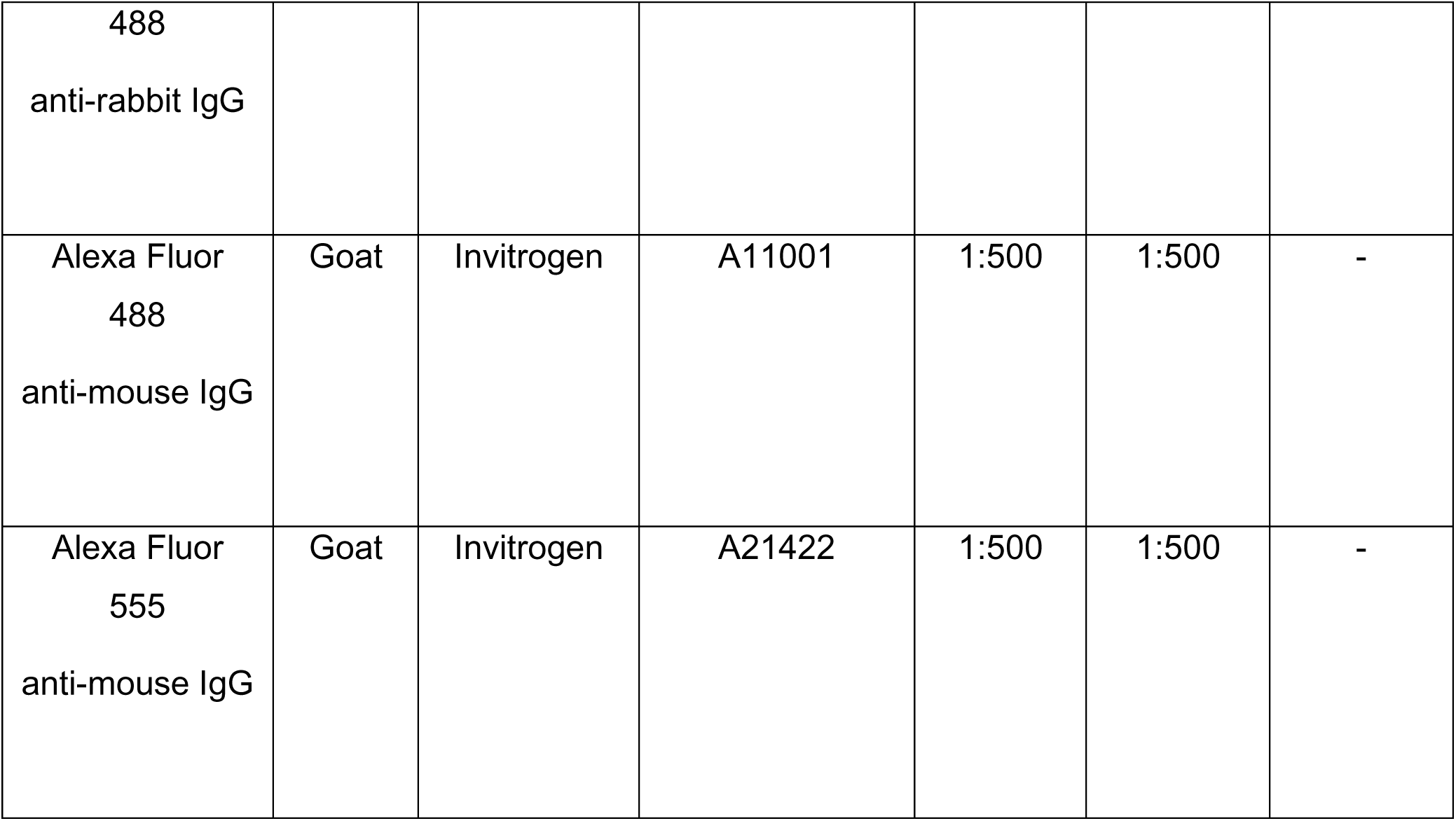
Antibodies used in this study.

### PARP1 recruitment to sites of laser irradiation

HeLa wild-type or HeLa POLE4 KO cells were grown in 8-well Lab-Tek II chambered coverglass 30 (Thermo Scientific) and transfected 48 h prior to imaging with GFP-tagged PARP1 chromobody (Chromotek). For sensitization, growth media was replaced with fresh medium containing 0.15 µg/mL Hoechst 33342 for 1 hour at 37°C. Prior to imaging, the sensitizing media was then replaced with CO_2_-independent imaging medium (Phenol Red-free Leibovitz’s L-15 medium (Life Technologies) supplemented with 20% fetal bovine serum, 2 mM glutamine, 100 µg/mL penicillin and 100 U/mL streptomycin). For PARP inhibition conditions, cells were treated with Olaparib (30 nM) for 30 min prior to imaging.

Live-cell imaging experiments were performed on a Ti-E inverted microscope from Nikon equipped with a CSU-X1 spinning-disk head from Yokogawa, a Plan APO 60x/1.4 N.A. oil-immersion objective lens and a sCMOS ORCA Flash 4.0 camera. Laser microirradiation at 405 nm was done along a 16 µm-line through the nucleus using a single-point scanning head (iLas2 from Roper Scientific) coupled to the epifluorescence backboard of the microscope. The laser power at 405 nm was measured prior to each experiment to ensure consistency across the experiments and set to 125 µW at the sample level. Cells were kept at 37°C with a heating chamber. Protein recruitment was quantified using a custom-made Matlab (MathWorks) routine.

### Immunofluorescence

For native BrdU staining, cells were grown with 20 µM BrdU-containing medium for 48 h, the media was then replaced with 10 µM Olaparib-containing medium for 24 h. Cells were washed with PBS, pre-extracted with 0.5% Triton X-100 in PBS for 5 min at 4°C then fixed with 4% paraformaldehyde (PFA) for 15 min at 4°C. Permeabilization was done using 0.5% Triton X-100 in PBS for 10 min followed by blocking with 5% FBS in 0.1% Triton X-100 for 45 min at room temperature, then incubated with primary antibody diluted in blocking solution overnight at 4°C.

For Rad51 experiments, cells were treated with 10 µM Olaparib-containing medium for the 48 h before being washed with PBS and pre-extracted with pre-extraction buffer (10 mM Tris-HCl, 2.5 mM MgCl2, 0.5% NP-40, 100× Protease inhibitor cocktail (Roche)) for 5 min at 4°C. Fixation was done using 4% PFA for 15 min at 4°C followed by permeabilization, blocking and antibody incubation as described earlier.

For S-phase PAR staining experiments, either wild-type or POLE4 KO cells were loaded with 2.5 μM of amine-reactive dye carboxyfluorescein diacetate, succinimidyl ester (CFSE) using the CellTrace™ CFSE Cell Proliferation Kit (Molecular Probes, Life Technologies) for 12 min at room temperature before seeding and mixing with the other unlabeled genotype. Cells were treated with DMSO (vehicle control) or PARGi (10 μM) or PARGi (10 μM) and Fen1i (10 μM) (Table 3) for 1h. Cells were pulse-labeled with the nucleotide analog EdU (10 μM) (5-ethynyl-2’-deoxyuridine, Baseclick, BCK-EdU555) for the last 20 min prior to fixing. Fixation and staining were done as described earlier.

**Table 3.** Inhibitors used in this study.

| Inhibitor | Commercial name | Company | Reference |
| --- | --- | --- | --- |
| AZD2281 | Olaparib | Selleck Chemicals | S1060 |
| AG014699 | Rucaparib | MedChem Express | HY-10617A |
| BMN-673 | Talazoparib | MedChem Express | HY-16106 |
| KU-55933 | ATMi | Selleck Chemicals | S1092 |
| VE-821 | ATRi | Selleck Chemicals | S8007 |
| NU7441 | DNAPKi | Selleck Chemicals | S2638 |
| PDD 00017273 | PARGi | MedChem Express | HY-108360 |
| LNT1 | Fen1i | Tocris | 6510 |

Following overnight incubation with the primary antibodies (Table 2), cells were washed three times with 0.1% Triton X-100 and incubated at room temperature with fluorescently tagged secondary antibody (Table 2) for 1 h. Next, cells were washed with 0.1% Triton X-100 and counterstained with DAPI (1 µg/mL in PBS) for 10 minutes. To detect proliferating cells, EdU incorporation was visualized by a Click-IT Kit (Baseclick) according to the manufacturer’s protocol.

Z-stacks of images were acquired on a Zeiss LSM800 confocal microscope with a Plan-Apochromat 20x/0.8 M27 or a water immersion Plan-Apochromat 40x/1.2 DIC M27 objective controlled by the ZEN 2.3 software. Fluorescence excitation was performed using diode lasers at 405, 488, 561 and 650 nm. Images were analyzed after generating maximum intensity projections of the z-stacks using a custom CellProfiler pipeline^78^.

### BrdU comet assay

Exponentially growing cells were plated in 24-well plate at a density of 3x10^5^ cells/well in duplicates to be left untreated or for Olaparib treatment in groteh media for 24 h. The following day the growth medium was changed to fresh DMEM containing 25 µM of the nucleotide analog ldU, the cells were incubated at 37°C for 30 min and then washed twice with PBS. After the labelling, the cells were washed two times with PBS and were cultured in Olaparib (20 µM) containing media for 24 h. Cells were harvested by trypsinization, collected and pelleted in DMEM washed with ice-cold PBS. Pelleted again and resuspended in 500 µL ice cold H_2_O_2_ (75 µM, diluted in PBS), kept on ice for 3 minutes and pelleted by centrifugation (5 min, 900 rpm at 4°C). H_2_O_2_ was removed by washing twice with ice-cold PBS and, after the last centrifugation, cells were resuspended in 70 µL 0.75% low melting agarose/slide. The following steps were performed as detailed in^44^ with slight modification detailed below. The alkaline lysis was performed in 0.3 M NaOH, 1 mM EDTA, pH 13 for 2 hours in Coplin jars. The DNA was left to unwind for 40 minutes in this ice-cooled electrophoresis buffer. The electrophoresis was subsequently conducted at 1 V/cm (25 V, 300 mA) for 30 minutes in the same buffer at 10 °C.

Following the electrophoresis, the slides were placed horizontally side by side on a glass tray and washed for 3x 5 min by gently layering with neutralization buffer (0.4 M Tris-HCl, pH 7.4). Slides were washed with two changes of PBS and blocked with PBS containing 1% BSA for 20 minutes at room temperature. The slides were incubated with 40 µL/gel of mouse monoclonal anti-BrdU (#347580, Becton Dickinson) (also reacts with IdU and CldU) in the dark in a humidified box at room temperature for two hours. The primary antibody was washed off with two changes of PBS and probed with 40 µL/gel of secondary antibody for 2 hours at room temperature in the dark. The slides were washed two times with PBS and air dried overnight in the dark at room temperature. Next day the slides were counterstained with propidium-iodide, mounted by Fluoromount mounting solution containing DAPI, covered with coverslips and stored at 4°C until microscopy. Imaging was performed using Zeiss Axioscope Z2 fluorescent microscope. Scanning of images was done using automated scanning platform of Metasystem and the quantitation of comets was done by Metasystems Neon Metafer4 software. Three independent experiments were done with duplicate slides, 150-300 comet/slides were scored and analyzed using GraphPad Prism7.

### Cell survival assays

POLE4 KO, POLE3 KO and their parental wild-type cells were seeded in defined numbers in 96-well plates and treated for one week with Olaparib (0, 0.45, 0.9, 1.8, 3.7, 7.5, 15, 30 µM), or Rucaparib (0, 0.45, 0.9, 1.8, 3.7, 7.5, 15 µM), or Talazoparib (0, 15, 31, 62, 125, 250 nM), or ATRi (0, 0.6, 1.2, 2.5, 5, 10 µM) (Table 3). For experiments with the combination of ATRi and Olaparib, one Olaparib concentration of (1 µM) was used along with an ATRi concentration of (0.6 µM). For experiments with RNAi-induced BRCA1 depletion the concentrations of Olaparib were (0, 0.3, 0.6, 1.2, 2.5, 5, 10 µM). Treatment with HU (Sigma-Aldrich, 0, 0.5, 1, 2, 4, 8 mM) was for 24h, with MMS (Sigma-Aldrich, 0, 0.0015, 0.003, 0.006, 0.012, 0.025, 0.05 %) for 1h, with Etoposide (Sigma-Aldrich, 0, 0.45, 0.9, 1.8, 3.7, 7.5, 15 µM) for 1h; following the indicated durations, cells were washed and incubated for one week in complete medium. After 7 days of incubation, the supernatants were aspirated and resazurin (Sigma) solution was added (25 µg/ml in Leibowitz’s L-15, Gibco). The fluorescent resorufin product was measured after 30-60 min using a Biotek Synergy H1 microplate reader with a 530/590 filter set.

### Flow cytometry for intracellular markers and cell cycle analysis

Cells were dissociated with TrypLe Select (Gibco), washed with PBS and fixed with ice-cold ethanol. For labelling the intracellular markers, the cells were permeabilized and blocked with 0.5% Triton X-100 and 5% FBS in PBS, and then incubated with primary antibody overnight against phospho-RPA2(T21) or phospho-H3(S10) at 4°C. Next, the cells were washed two times with PBS, and incubated with fluorescently tagged secondary antibodies for 2 h at room temperature. Finally, the DNA staining solution was added (10 μg/mL propidium-iodide and 10 μg/mL RNase in PBS) for 15 min at room temperature. The samples were analyzed with CytoFLEX S flow cytometer (Beckman Coulter Life Sciences) or FACSCalibur (Becton Dickinson). The measurements were evaluated with Kaluza Analysis software (Beckman Coulter Life Sciences).

### PARylation assay

Cells were cultured in 6 cm dishes. The cells were treated with H_2_O_2_ (2 mM) in fresh culturing medium for the indicated timepoints. At the time of collection, the cells were washed twice in 1X PBS and lysed directly using denaturing lysis buffer (4% SDS, 50 mM Tris-HCl, pH 7.4, 100 mM NaCl, 4 mM MgCl_2_, 5 U/µl Benzonase). The cell lysates were collected using a cell scraper and the total protein concentration was equalized after measuring the initial concentration by NanoDrop (A280 setting). Samples were boiled in 4x Laemmli buffer for 5 min at 95°C prior to western blotting.

### Western blotting

Protein samples were prepared for SDS–polyacrylamide gel electrophoresis in 4× sample buffer (10% SDS, 300 mM Tris-HCl, 10 mM β-mercaptoethanol, 50% glycine, and 0.02% bromophenol blue). Separated proteins were blotted onto nitrocellulose or PVDF membranes, blocked for 1 hour at RT in 5% low-fat milk or 5% BSA in 0.1% Tris-buffered saline, and incubated with primary antibodies overnight at 4°C. For secondary antibodies, horseradish peroxidase-conjugated secondary antibodies were used for 1 hour. Membranes were developed with enhanced chemiluminescence using Odyssey Fc Imaging System (LI-COR Biotechnology).

### Statistical analysis

All experiments were done at least in triplicate and for immunofluorescence experiments at least 200 cells were scored. A minimum of 10 cells were irradiated in live-cell imaging experiments. Graphing and statistical analysis were done using GraphPad Prism versions 6 and 7. Statistical analysis of cell survival experiments was done using two-way ANOVA. PARP1 recruitment experiments were analyzed using Mann-Whitney unpaired t-test. Statistics for immunofluorescence experiments were performed using one-way ANOVA. Asterisks represent *p* values, which correspond to the significance (\**p* < 0.05, \*\**p* < 0.01, \*\*\**p* < 0.001 and \*\*\*\**p* < 0.0001)

### Data Availability

The data that support the findings of this study are available from the corresponding author upon reasonable request.

## 5 ACKNOWLEDGEMENTS

We would like to thank the technical assistance of Adrián Kószó in the laboratory of G.T. and that of the Microscopy Rennes Imaging Center (BIOSIT, Université Rennes 1). The work in the Timinszky laboratory was supported by the National Research Development and Innovation Office (K143248). A.G.K. was supported by the National Academy of Scientist Education Program of the National Biomedical Foundation under the sponsorship of the Hungarian Ministry of Culture and Innovation and the New National Excellence Program of the Hungarian Ministry of Culture and Innovation (UNKP-22-3-SZTE-264). For this work, S.H.’s group received financial support from the Agence Nationale de la Recherche (AROSE, ANR-22-CE12-0039), the Institut National du Cancer (PLBIO-2019) and the Institut Universitaire de France. Research in the Haracska laboratory was supported by the National Research, Development and Innovation Office (PharmaLab, RRF-2.3.1-21-2022-00015 and TKP-31-8/PALY-2021). The models in Figure 6 were created with BioRender.

## AUTHORS CONTRIBUTIONS

G.T. conceived the project. H.M., R.F.B. and G.T. planned the research. H.M. generated POLE3 and POLE4 knockouts. H.M., R.F.B., M.M., E.P.J. and A.G.K. did cell proliferation assays and immunofluorescence microscopy. R.F.B. performed flow cytometry assays. S.Z., R.S. and S.H. performed live cell imaging assays. A.M. performed H_2_O_2_ treatment and western blotting. M.M. and L.H. performed comet assays. H.M drafted the manuscript. R.F.B and G.T. reviewed and edited the manuscript. All authors read and commented on the manuscript.

