## Supplemental Figures for "The loss of DNA polymerase epsilon accessory subunits POLE3-POLE4 leads to BRCA1-independent PARP inhibitor sensitivity"

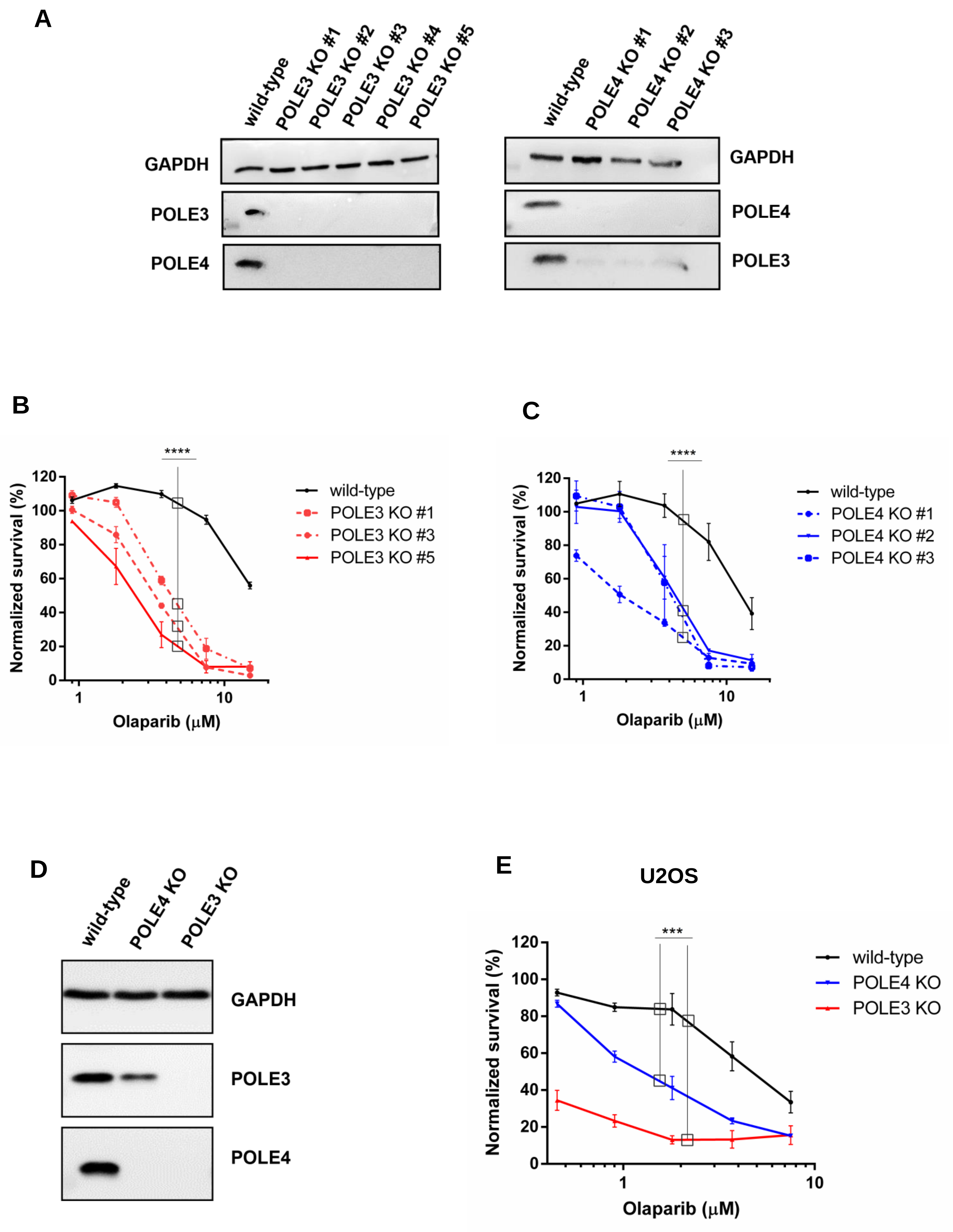


Supplementary Figure 1

Mamar et al.

**Supplementary Figure 1:**

(A) Western blot of different independent clones of (left) POLE3 KO and (right) POLE4 KO and their parental HeLa wild-type. GAPDH is used as a loading control.

(B, C) Cell survival assays demonstrating sensitivity of (B) POLE3 KO clones and (C) POLE4 KO clones to Olaparib treatment compared to their parental HeLa wild-type. PARPi treatment was refreshed once during the 7-day long experiment. Mean ± SEM (n=3). Asterisks indicate *p*-values obtained by two-way ANOVA (**** $p < 0.0001$).

(D) Western blot of POLE3 and POLE4 knockouts generated in the U2OS cell line showing the importance of the accessory subunits for their stability. GAPDH is used as a loading control.

(E) Cell survival assays of U2OS wild-type, POLE3 and POLE4 knockout cells. The treatment lasted for 7 days and was refreshed once. Mean ± SEM (n=3). Asterisks indicate *p*-values obtained by two-way ANOVA (*** $p < 0.001$).

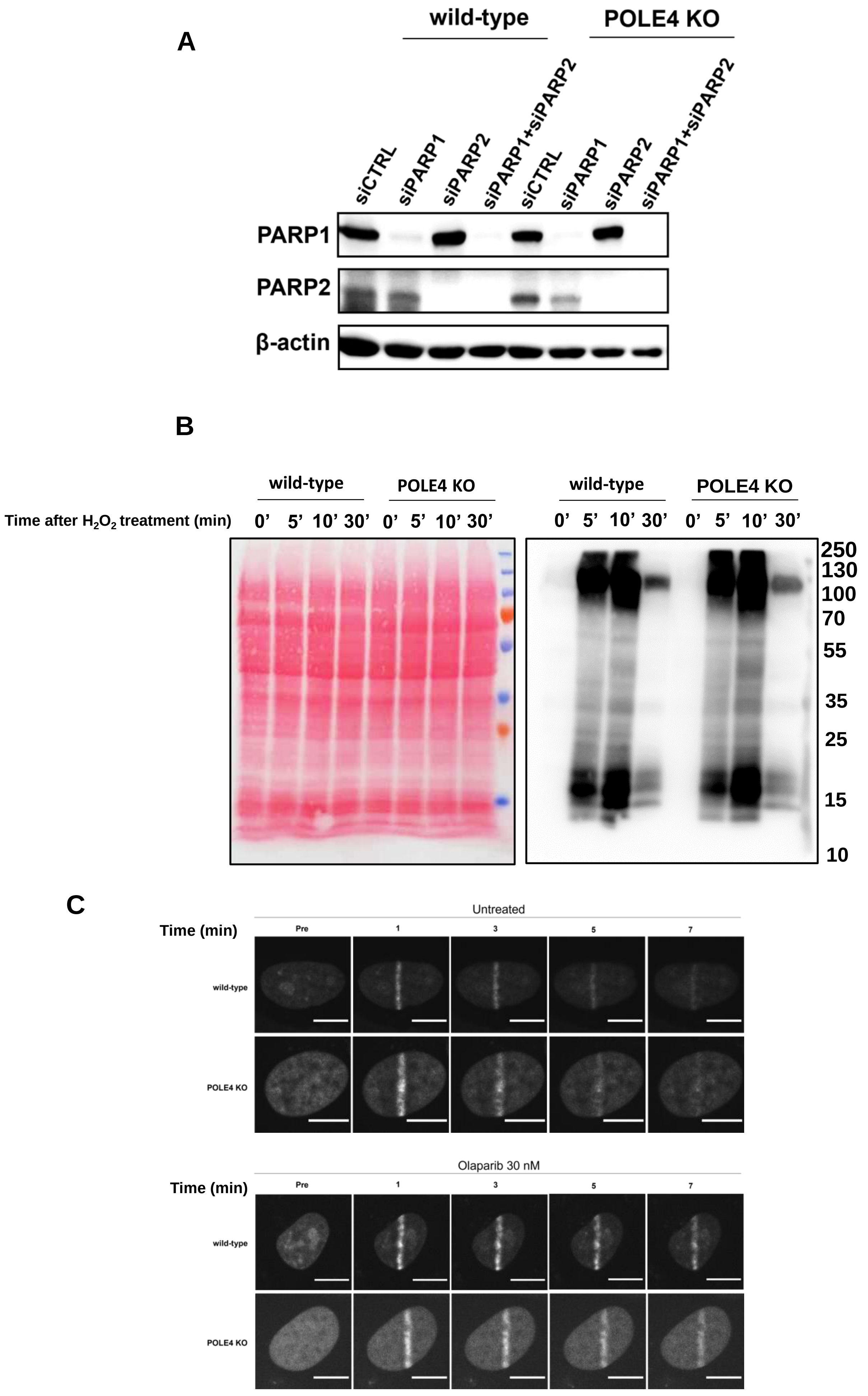
A
wild-type
POLE4 KO
siCTRL
siPARP1
siPARP2
siPARP1+siPARP2
siCTRL
siPARP1
siPARP2
siPARP1+siPARP2
PARP1
PARP2
β-actin
B
wild-type
POLE4 KO
wild-type
POLE4 KO
Time after $H_2O_2$ treatment (min)
0’ 5’ 10’ 30’ 0’ 5’ 10’ 30’
0’ 5’ 10’ 30’ 0’ 5’ 10’ 30’
250
130
100
70
55
35
25
15
10
C
Untreated
Time (min)
Pre
1
3
5
7
wild-type
POLE4 KO
Olaparib 30 nM
Time (min)
Pre
1
3
5
7
wild-type
POLE4 KO

**Supplementary Figure 2:**

(A) Western blot of HeLa wild-type and POLE4 KO cells showing downregulation of PARP1, PARP2, or both due to transfection with the indicated siRNA. The cells were harvested for western blot 48h post transfection. β-Actin is used as a loading control.

(B) ADPr levels in both Hela wild-type and POLE4 KO shown by western blotting. The cells were treated or not with $H_2O_2$ (2 mM) for the indicated timepoints. ADPr signal was probed using anti-pan-ADPr antibody. Ponceau staining is used as a loading control.

(C) Representative images of GFP-tagged PARP1 chromobody accumulation at sites of laser-induced damage in HeLa wild-type and POLE4 KO cells treated or not with Olaparib (30 nM). Scale bar, 10 μm.

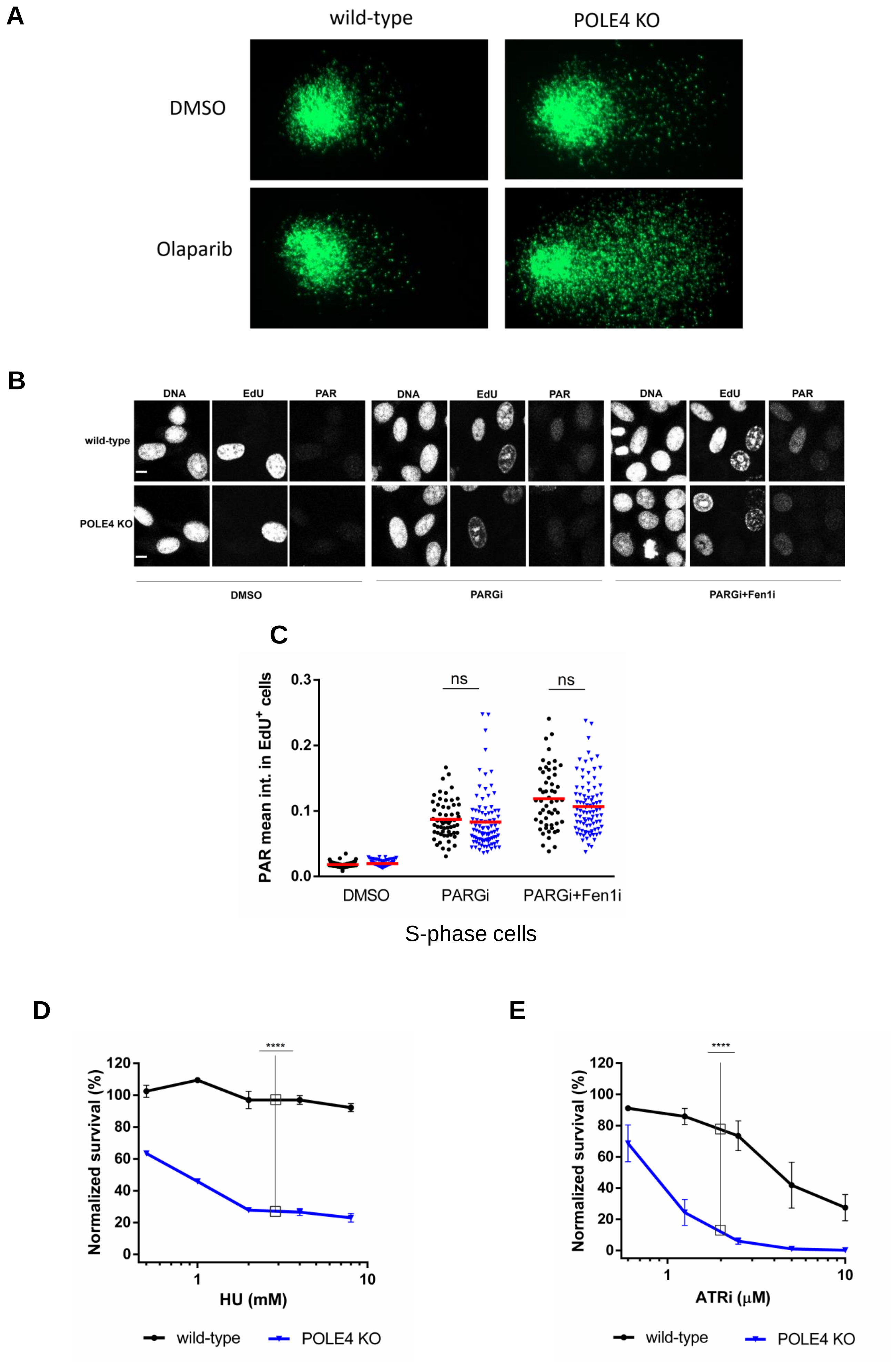


**Supplementary Figure 3**

**Supplementary Figure 3:**

(A) Representative images of BrdU comet experiment showing tail formation in POLE4 KO compared to HeLa wild-type upon treatment of Olaparib (20 μM, 24h).

(B) Representative images of HeLa wild-type and POLE4 KO cells showing PAR signal in EdU-positive cells (indicative of S-phase) upon treatment with PARGi (10 μM, 1h) and PARGi (10 μM) + Fen1i (10 μM) for 1h. Scale bar, 10 μm.

(C) Immunofluorescence experiment of HeLa wild-type and POLE4 KO cells showing no significant difference in the mean intensity of PAR signal in S-phase cells between the two compared genotypes. Cells were treated or not with PARGi (10 μM) or PARGi (10 μM) + Fen1i (10 μM) for 1h, and with EdU (10 μM, last 20 min) before fixation. PAR signal was detected using anti-PAR WWE reagent (Millipore). EdU click-it reaction was used to identify cells in S-phase. The graphs are derived from a representative experiment out of three independent repetitions. Statistical analysis is done using one-way ANOVA (ns: not significant).

(D) Cell survival assay showing POLE4 KO sensitivity to hydroxyurea (HU) treatment compared to HeLa wild-type. HU treatment was for 24h, then the cells were left to recover for 7 days in culturing media. Mean ± SEM (n=3). Asterisks indicate *p*-values obtained by two-way ANOVA (**** $p < 0.0001$).

(E) Cell survival assay of HeLa wild-type and POLE4 KO cells upon treatment with the indicated concentrations of ATRi. The treatment lasted for 7 days and was refreshed once. Mean ± SEM (n=3). Asterisks indicate *p*-values obtained by two-way ANOVA (**** $p < 0.0001$).

**A**

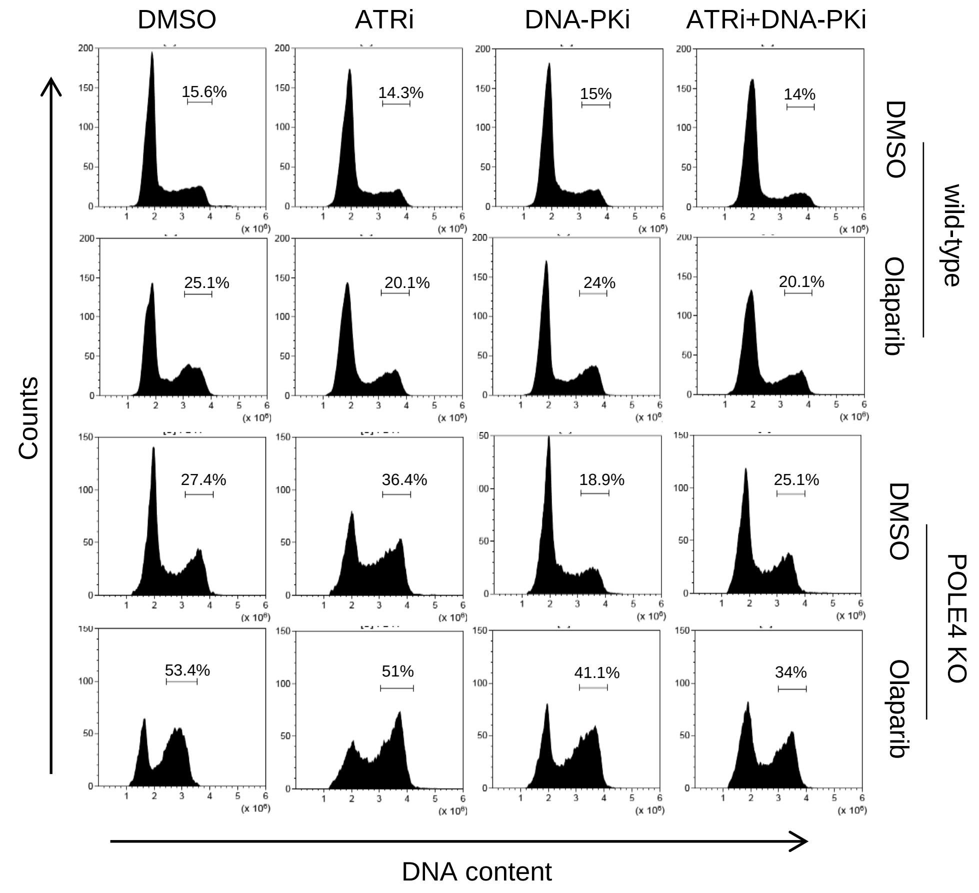


**B**

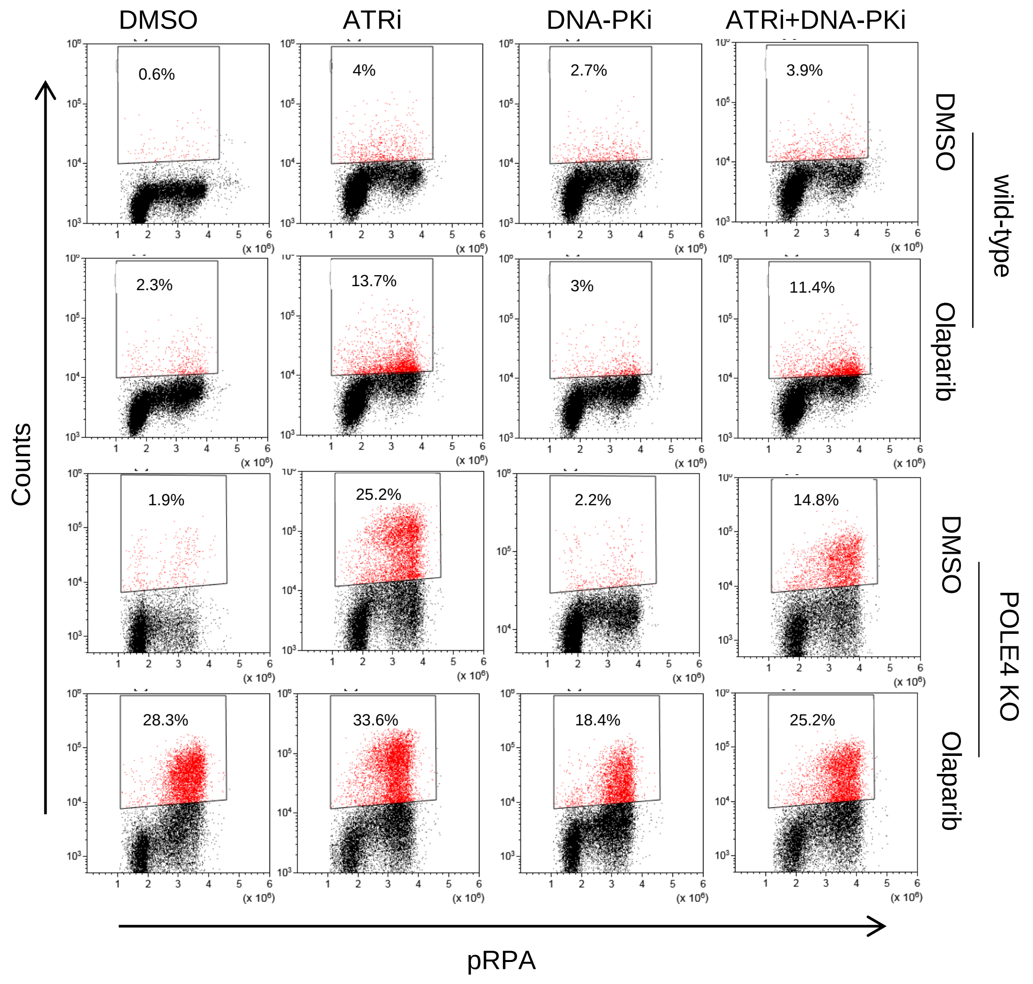


**Supplementary Figure 4** **Mamar et al.**

**Supplementary Figure 4:**

(A) Flow cytometry experiment showing cell-cycle profile of Hela wild-type and POLE4 KO cells after 24h treatment with Olaparib (5 μM) and/or ATRi (5 μM), DNA-PKi (5 μM) or both. DMSO was used as a solvent control. The cells were fixed and stained with propidium-iodide (DNA content). Numbers represent the percentages of G2/M population relative to all cells. The figure is a representative of three independent experiments.

(B) Flow cytometry of Hela wild-type and POLE4 KO cells after 24h treatment of Olaparib (5 μM) and/or DNA-PKi (5 μM), ATRi (5 μM) or both. DMSO was used as a solvent control. The cells were fixed and stained with anti-pRPA and propidium-iodide (DNA content). Percentages of pRPA positive cells (red) are shown. The figure is a representative of three independent experiments.

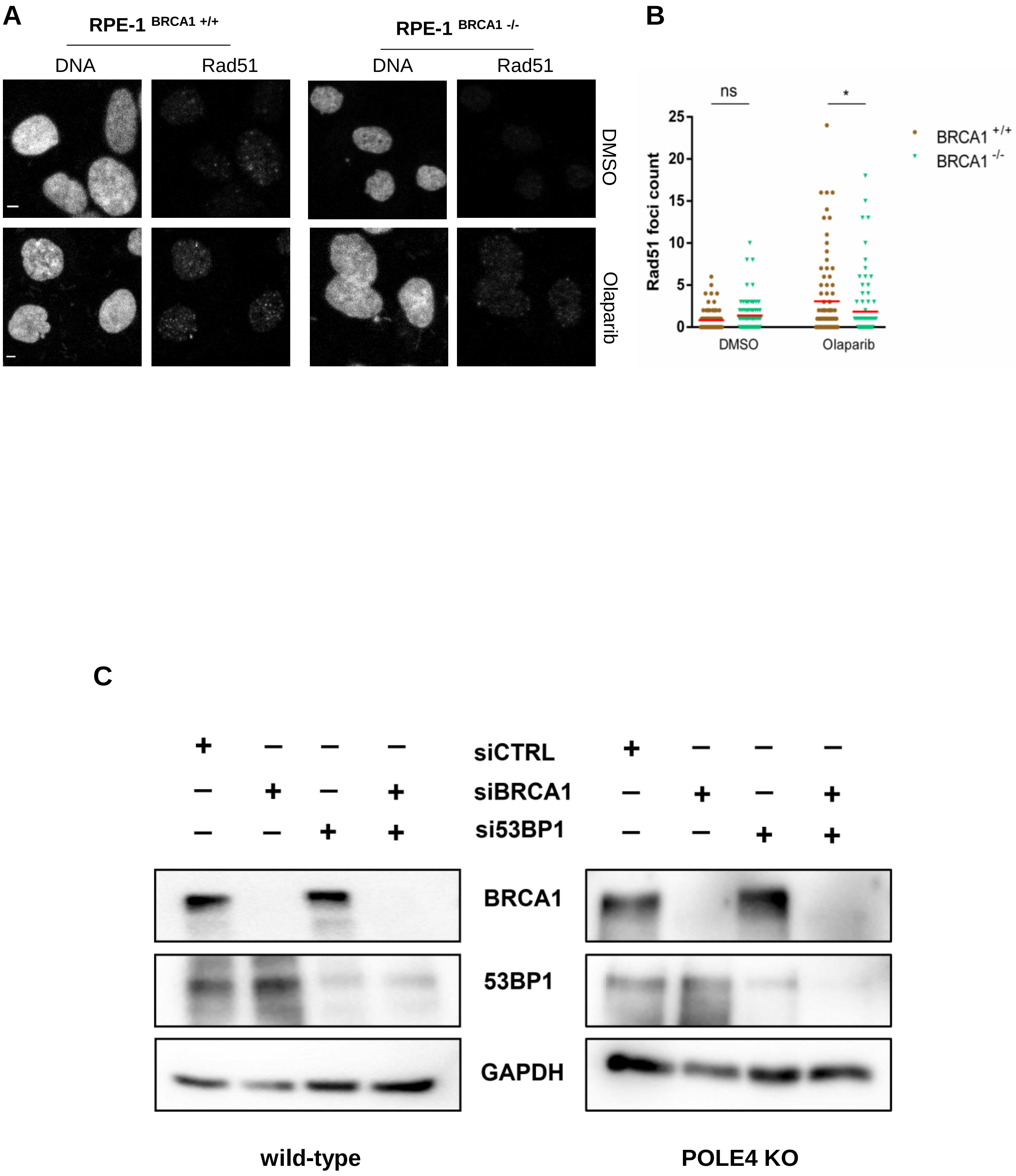


Supplementary Figure 5

Mamar et al.

**Supplementary Figure 5:**

(A) Immunofluorescence experiment of Rad51 foci formation in RPE-1 BRCA1-deficient cells and their wild-type upon treatment with Olaparib (10 μM, 48h), Scale bar, 10 μm.

(B) Quantification of Rad51 foci count in RPE-1 BRCA1-deficient cells and their wild-type upon treatment with Olaparib (10 μM, 48h). The experiment is representative of three independent repetitions. Asterisks indicate *p*-values obtained by one-way ANOVA (ns. Not significant, * $p < 0.05$).

(C) Representative western blot of HeLa wild-type and POLE4 KO cells demonstrating depletion of the indicated proteins 48h post treatment with the indicated siRNA. GAPDH is used as a loading control.
